# PURA Syndrome-causing mutations impair PUR-domain integrity and affect P-body association

**DOI:** 10.1101/2023.09.19.558386

**Authors:** Marcel Proske, Robert Janowski, Sabrina Bacher, Hyun-Seo Kang, Thomas Monecke, Tony Köhler, Saskia Hutten, Jana Tretter, Anna Crois, Lena Molitor, Alejandro Varela-Rial, Roberto Fino, Elisa Donati, Gianni De Fabritiis, Dorothee Dormann, Michael Sattler, Dierk Niessing

## Abstract

Mutations in the human *PURA* gene cause the neuro-developmental PURA syndrome. In contrast to several other mono-genetic disorders, almost all reported mutations in this nucleic acid binding protein result in the full disease penetrance. In this study, we observed that patient mutations across PURA impair its previously reported co-localization with processing bodies. These mutations either destroyed the folding integrity, RNA binding or dimerization of PURA. We also solved the crystal structures of the N- and C-terminal PUR domains of human PURA and combined them with molecular dynamics simulations and NMR measurements. The observed unusually high dynamics and structural promiscuity of PURA indicated that this protein is particularly susceptible to mutations impairing its structural integrity. It offers an explanation why even conservative mutations across PURA result in the full penetrance of symptoms in patients with PURA syndrome.

## INTRODUCTION

The family of purine-rich element binding (PUR) protein is highly conserved from plants to humans (Molitor *et al*, 2021). Amongst the three vertebrate paralogs PURA, PURB, and PURG, the best-studied family member is PURA (Bergemann & Johnson, 1992; Haas *et al*, 1993; Haas *et al*, 1995). It is ubiquitously expressed and has been implicated in several cellular processes, including transcriptional and translational gene regulation as well as mRNA transport in neurons (Chepenik *et al*, 1998; Gallia *et al*, 2001; Haas *et al*., 1993; Haas *et al*., 1995; Johnson *et al*, 2006; Kobayashi *et al*, 2000; Mitsumori *et al*, 2017; Tretiakova *et al*, 1999). Results from two independent knock-out mouse models demonstrated that PURA is important for postnatal brain development (Hokkanen *et al*, 2012; Khalili *et al*, 2003).

Accordingly, PURA has been implicated as a modulator of neurodegenerative disorders, such as fragile X-associated tremor/ataxia syndrome (FXTAS), and the amyotrophic lateral sclerosis (ALS) frontotemporal dementia (FTD) spectrum disorder (Swinnen *et al*, 2020). PURA was reported to be present in the pathological RNA foci of both disorders and to protect against RNA toxicity upon ectopic overexpression (Mori *et al*, 2013; Rossi *et al*, 2015; Shen *et al*, 2018; Swinnen *et al*, 2018; Xu *et al*, 2013). In addition, PURA was shown to co-localize with an ALS-causing mutant variant of the FUS protein in stress granules of ALS patients. Importantly, overexpression of PURA reduced the toxicity of mutant FUS by preventing its mis-localization (Daigle *et al*, 2016).

In 2014, two studies reported that a monogenetic neurodevelopmental disorder is caused by sporadic mutations in the *PURA* gene (Hunt *et al*, 2014; Lalani *et al*, 2014). Hallmarks of this so-called PURA syndrome are developmental delay, moderate to severe intellectual disability, hypotonia, epileptic seizures, and feeding difficulties, amongst others (Reijnders *et al*, 2018) (Johannesen *et al*, 2021). Mutations causing PURA syndrome are often frame-shift events but also point mutations distributed over the entire sequence.

Based on the crystal structures of PURA from *Drosophila melanogaster* (*dm*PURA), three conserved sequence regions termed PUR repeats I, II, and III were identified to fold into globular domains (Graebsch *et al*, 2010; Graebsch *et al*, 2009; Weber *et al*, 2016). Whereas PUR repeat I and II assemble into an N-terminal PUR domain via intramolecular interactions, two C-terminal PUR repeats III from different molecules interact with each other and thus mediate dimerization of PURA. All these PUR domains belong to the class of the PC4-like protein family, which mainly bind single-stranded nucleic acids and can unwind double-stranded DNA and RNA (Janowski & Niessing, 2020). Although both PUR domains possess the typical PC4-like β-β-β-β-α-(linker)-β-β-β-β-α topology, the N-terminal domain has been suggested to be the main RNA/DNA interaction entity (Weber *et al*., 2016). In a previous study, homology models based on the *dm*PURA crystal structures (Graebsch *et al*., 2009; Weber *et al*., 2016) were used to predict *in silico* the effects of PURA syndrome-causing mutations on the structural integrity of the human PURA protein (Reijnders *et al*., 2018). While these mutations could be classified into groups that likely either do or do not impair the structural integrity of PURA, these predictions remained speculative (Reijnders *et al*., 2018). Surprisingly, in contrast to phenotypically related genetic diseases such as the Rett syndrome (Lombardi *et al*, 2015), no hot-spot regions could be identified in the protein sequence that trigger the PURA syndrome. With the exception of the unstructured N-terminal region and the very C-terminus, almost all mutations across the protein sequence appear to result in the full disease spectrum (Johannesen *et al*., 2021; Reijnders *et al*., 2018). To date, we fail to understand why there is such an underrepresentation of mild phenotypes in patients with PURA syndrome.

In this study, we experimentally assessed the structure and function of human PURA as well as the impact of representative PURA syndrome-causing mutations on the protein’s integrity. When studying the impact of patient mutations on PURA’s subcellular localization, we observed impaired processing body (P-body) association but normal localization to stress granules, indicating a potential importance of P-bodies for PURA-syndrome pathology. Two particularly interesting disease-causing mutations, K97E and R140P, were further analyzed by structural means. The results indicated that the N-terminal PUR domain is unusually flexible in its fold and prone to misfolding upon mutation. In summary, we provide a structure-based explanation for the underrepresentation of mutations with mild symptoms in patients and a rational approach for predicting and functionally testing mutations of uncertain significance.

## RESULTS

### Effect of the mutations in *hs*PURA on stress granules association

Several patient-related mutations are frame-shift events and thus predictably destroy the integrity of PURA. In addition, a number of point and indel mutations have been reported for which the pathology-causing effects are less easy to predict (Johannesen *et al*., 2021; Reijnders *et al*., 2018). To understand the functional impact of such mutations, we decided to first study the effect of three known patient mutations, K97E, I206F, and F233del (Hunt *et al*., 2014; Lalani *et al*., 2014; Reijnders *et al*., 2018) (**Fig 1A**), on the subcellular localization of *hs*PURA.

**Figure 1.**
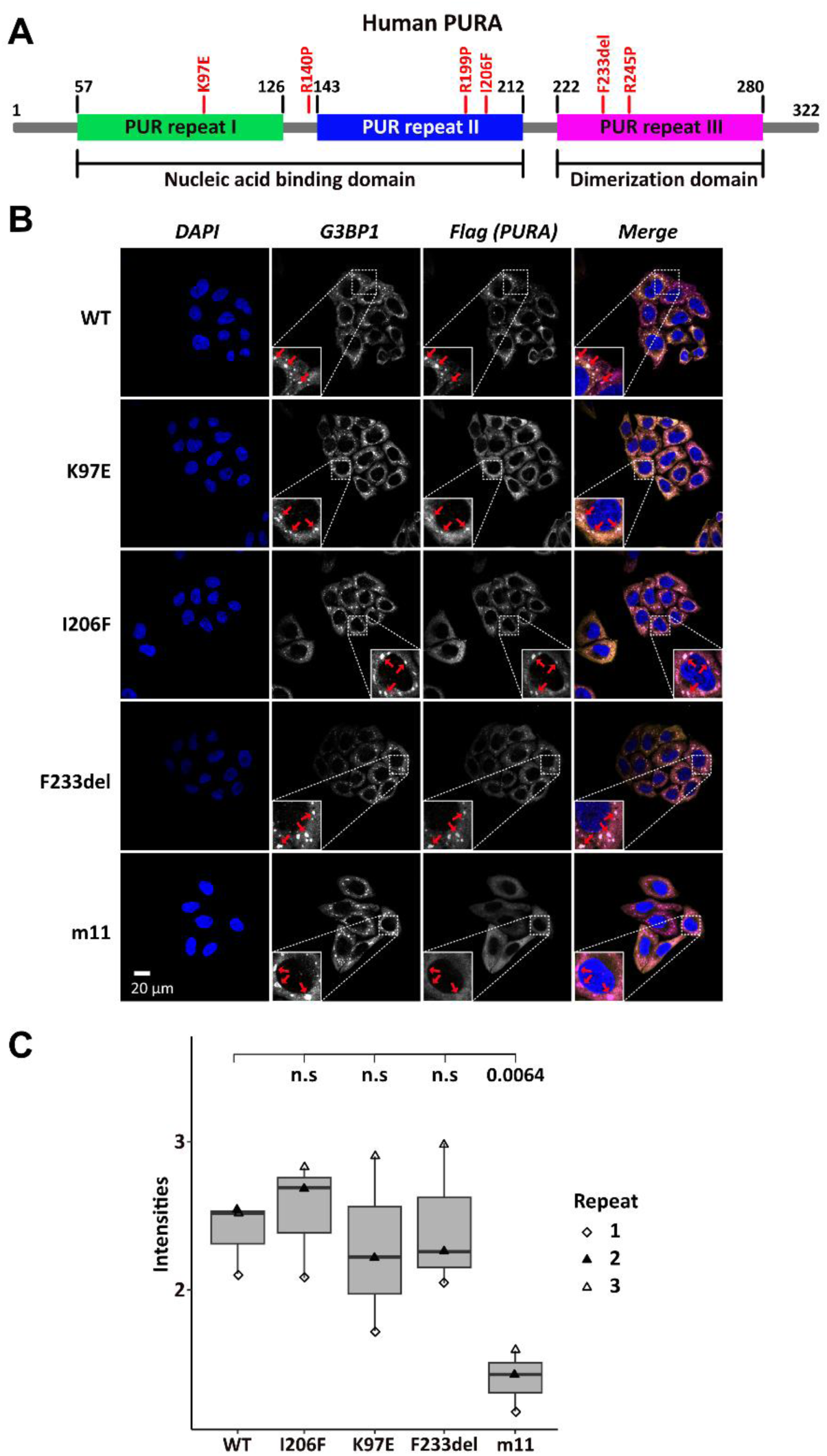
Stress granule localization of *hs*PURA depends on its ability to bind RNA. **A** Schematic overview of the hsPURA protein. Patient-derived mutations experimentally assessed in this study are marked in red. **B** In HeLa cells stress granules were identified by G3BP1 immunostaining (magenta) and overexpressed *hs*PURA (yellow) by its N-terminal Flag-tag. Nuclei were stained with DAPI (blue). Representative stress granules are marked with red arrows in the zoomed-in area. Except for *hs*PURA m11, all overexpressed versions of *hs*PURA (wild-type and mutant) showed strong stress-granule localization. **C** Box-Whiskers plot of quantified intensities of *hs*PURA staining in stress granules above cytoplasmic background levels upon arsenite-induced stress treatment (see A). Three biological replicates (indicated by different icons) with 66 stress granules each were quantified for each cell line. Box-Whiskers-Plot of average signal intensity per repeat in each cell line. No significant difference was detected between patient-related mutations and WT. Only for the m11 mutant protein the association to stress granules was significantly reduced (p=0.0064), indicating the requirement of RNA binding for stress-granule association. P-value were calculated using unpaired two-sided students t.test of mutant compared to WT Ctrl.

Previous studies in human cell cultures indicated that *hs*PURA localizes to stress granules upon cellular stress and may be essential for their formation (Daigle *et al*., 2016; Markmiller *et al*, 2018). To analyze whether the afore-mentioned patient-related mutations affect the localization of *hs*PURA to stress granules, the mutant and wild-type versions of full-length FLAG-tagged *hs*PURA were overexpressed in HeLa cells using a doxycycline-inducible expression cassette and cells were stressed for one hour with 500 µM sodium arsenite (**Appendix Fig S1** and **Fig 1B**). To find out whether stress-granule localization of *hs*PURA depends on its nucleic-acids binding properties, we also utilized an RNA-binding deficient version of *hs*PURA bearing eleven structure-guided point mutations (m11) as a control (for rationale and design of this mutant, see **Appendix Fig S2**). Using an anti-FLAG tag antibody, we observed wild-type *hs*PURA to be distributed within the cytoplasm and accumulated in stress-granule structures, as seen by co-staining with the stress-granule marker G3BP1 (**Fig 1B**). In contrast, the nucleic acid-binding deficient mutant *hs*PURA m11 failed to co-localize to G3BP1-positive granules, indicating that RNA-binding by *hs*PURA is necessary for its co-localization with stress granules. Furthermore, all three versions of *hs*PURA bearing patient-derived mutations localized to stress granules upon cellular stress (**Fig 1B**). In all three patient-related mutations, no significant reduction of PURAs stress granule association was seen when compared to the wild-type control (Fig 1C).

To assess the previously reported influence of *hs*PURA on stress-granule formation at endogenous PURA levels (Daigle *et al*., 2016), we performed an siRNA-mediated knockdown of *hs*PURA in HeLa cells and stained with the stress-granule marker G3BP1 and PURA. Efficient knockdown of endogenous PURA levels was validated with western blot experiments, showing a significant reduction of PURA protein upon knockdown (S3A-C). As expected, endogenous *hs*PURA co-localized with G3BP1 upon stress treatment with 500 µM arsenite PURA (**Appendix Fig S4A** and **S4B**) and thus behaved similarly to overexpressed *hs*PURA. A PURA knockdown did not alter the total number of microscopically visible stress granules but resulted in a greater ratio of smaller stress granules (**Appendix Fig S4C**). Although this finding suggests that PURA is not essential for stress granule formation, it should be noted that the knockdown did not completely eliminate PURA levels and hence a stronger effect on stress granules could occur in a complete knockout of PURA. However, PURA-syndrome patients bear a heterozygous mutation, suggesting that an incomplete knockdown likely resembles the pathogenic situation more closely than a full knockout. Together these findings suggest that (i) reduced stress-granule association of mutant *hs*PURA variants may not be a core feature of the PURA syndrome and (ii) impaired stress-granule formation is not very likely to be directly responsible for the etiology of this disorder.

### Localization of *hs*PURA to P-bodies is impaired by PURA syndrome-causing mutations

To assess the localization of *hs*PURA to cytoplasmic granules in unstressed conditions, we first recapitulated its recently reported localization to P-bodies (Molitor *et al*, 2023). Co-staining with the marker DCP1A indeed confirmed a co-localization of overexpressed *hs*PURA in P-bodies of HeLa cells (**Fig 2A** and **2B**). As for stress granules, the PURA m11 mutant failed to accumulate in P-bodies, indicating that RNA binding of PURA contributes to its P-body localization. In addition, the K97E and F233del mutants of *hs*PURA showed significantly reduced co-localization to P-bodies (**Fig 2A** and **2B**). In summary, these experiments indicate that P-body localization of *hs*PURA requires its RNA-binding activity, and that the majority of tested patient mutations result in impaired P-body localization. Furthermore, since F233del mutant has been predicted to have impaired dimerization properties (Reijnders *et al*., 2018), these findings suggest dimerization to be important for PURA’s P-body association.

**Figure 2.**
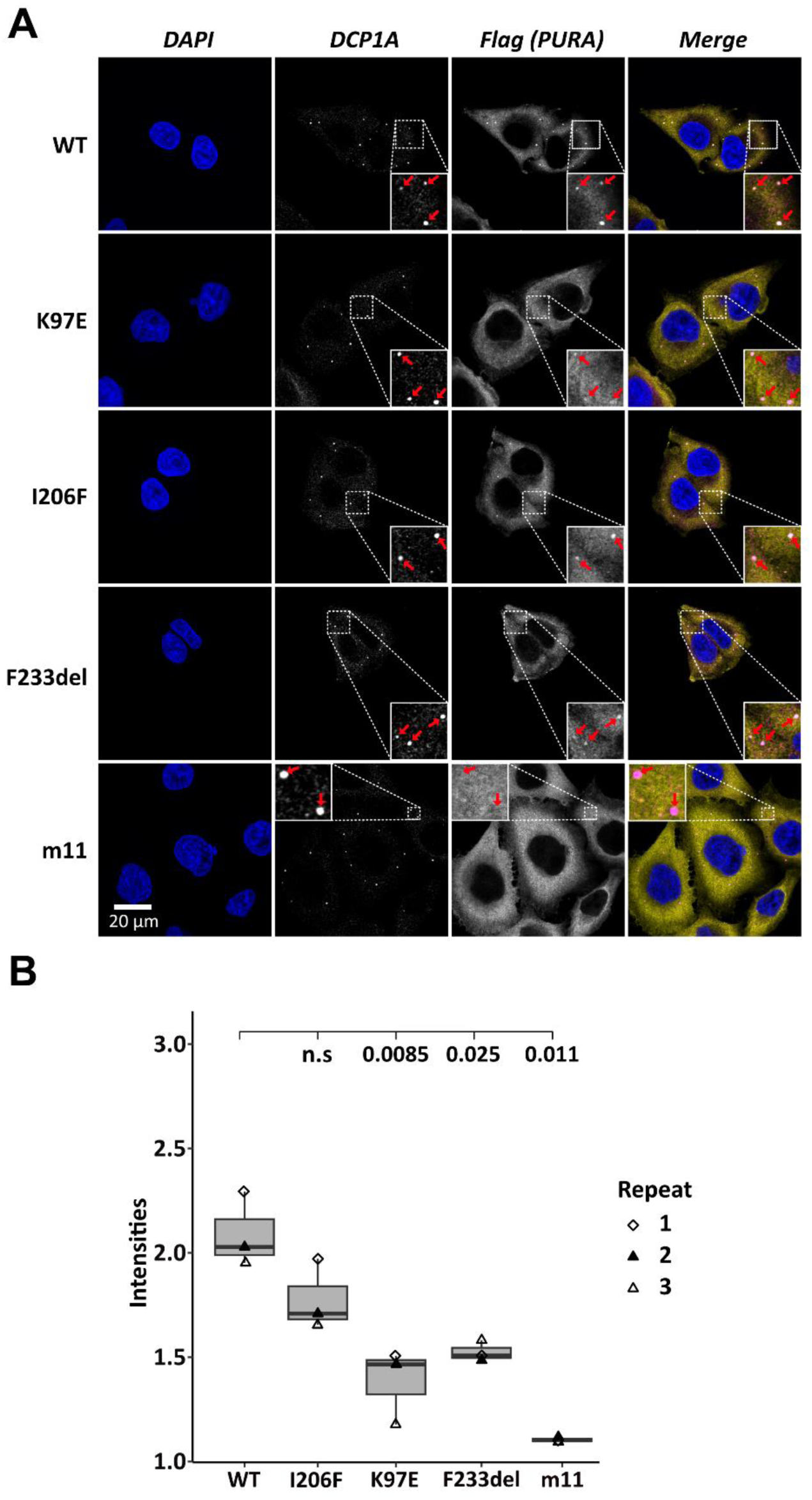
Processing-body association of *hs*PURA in HeLa cells is impaired by patient-derived mutations. **A** P-bodies were identified by immunostaining against DCP1A and against the N-terminal FLAG-tag of overexpressed *hs*PURA FL. Whereas their individual staining is shown in white, the overlay of DCP1A and FLAG is shown in magenta and yellow, respectively. Nuclei were stained with DAPI (blue). All cell lines show P-body formation, marked with red arrows in the zoomed area. **B** Box-Whiskers plots of quantification of intensities of *hs*PURA staining in P-bodies above cytoplasmic background levels. For each cell line, three biological replicates (indicated by different icons) with each 100 P-bodies were quantified, and Box-Whiskers-Plot generated for each cell line. Significantly lower P-body localization was observed for mutant K97E (p = 0.0085), F2333del (P = 0.025) and m11 (p =0.011) variants of *hs*PURA. P-value were calculated using unpaired two-sided students t.test of mutant compared to WT Ctrl.

### *hs*PURA does not readily undergo RNA-driven phase separation *in vitro*

*hs*PURA accumulates in an RNA-dependent fashion to stress granules as well as to P-bodies. Both types of granules are liquid-phase separated entities, indicating that the recruitment to such granules might occur via phase separation of RNA-bound *hs*PURA. Since *hs*PURA was recently shown to be required for P-body formation in HeLa cells and fibroblasts (Molitor *et al*., 2023), PURA-dependent liquid phase separation could potentially also directly contribute to the formation of these granules. Furthermore, a loss of such a function could potential contribute to the etiology of PURA syndrome. In order to test this hypothesis, we used a microscope-based approach to visualize the ability of RNA-bound *hs*PURA to phase separate *in vitro*. As a positive control for RNA-mediated liquid-phase separation, the protein FUS RGG3 was used (Hofweber *et al*, 2018). In contrast to this control, no indication of phase separation was observed for *hs*PURA either in the presence or absence of total HeLa cell RNA (**Appendix Fig S5**). These results indicate that *hs*PURA does not readily phase separate with total cellular RNA under the tested experimental conditions. Of note, this observation does not exclude the possibility that *hs*PURA can undergo phase separation under different experimental conditions, for instance in presence of additional co-factors. However, when putting this observation in perspective with previous reports, it seems unlikely that P-body formation directly depends on phase separation by *hs*PURA, but rather on its recently reported function as gene regulator of the essential P-body core factors LSM14a and DDX6 (Molitor *et al*., 2023).

### Recombinant expression tests of PURA protein with disease-causing mutations

To understand the impact of mutations on the molecular and structural integrity of *hs*PURA, we recombinantly expressed the above-described mutant *hs*PURA proteins K97E, I206F, and F233del in *E. coli*. In addition, *hs*PURA with either of the mutations R140P, R199P, and R245P were recombinantly expressed (**Fig 1A**). All three mutations exchange an arginine against a proline, albeit in different positions of the protein and very different structural contexts. R140P was initially reported to be a disease-causing mutation (Lee *et al*, 2018) but later re-interpreted as a mutation for which it is unclear if it can trigger PURA Syndrome, i.e. of uncertain significance (https://www.ncbi.nlm.nih.gov/clinvar/variation/192343/). In contrast, the mutations R199P and R245P have been described as disease-causing, with matching symptoms for patients with PURA syndrome (Reijnders *et al*., 2018). Hence, R140P was chosen in this study to explore the diagnostic potential of our approach, whereas R199P and R245P served as a reference point for pathology-inducing mutations.

Even after extensive optimization, recombinantly expressed *hs*PURA fragments with the mutations R199P and I206F (full-length variant and N-terminal PUR domain consisting of repeats I-II; PURA I-II) as well as F233del and R245P (full-length variant and C-terminal PUR domain consisting of two repeats III; PURA III) could not be purified due to protein insolubility (**Appendix Table S1**). In contrast, N-terminal PUR domains (PURA I-II) with either of the two remaining mutations K97E and R140P were soluble and could be purified to near homogeneity. The observed solubility of recombinant *hs*PURA I-II K97E is also in agreement with the previous *in silico* prediction (Reijnders *et al*., 2018) as K97 has a surface-exposed side chain that is unlikely to cause folding issues upon mutation into another flexible, polar side chain. For the R140P mutation with unclear pathological significance, no conclusive prediction on its impact on protein folding could be made in the past.

### The C-terminal dimerization PUR domain adopt a classical PC4-like fold

Several PURA-Syndrome-causing mutations have been reported to be located in the C-terminal, dimerization-mediating PUR domain (Johannesen *et al*., 2021; Reijnders *et al*., 2018). Due to the lack of an experimental structure of human PURA, previous predictions of the impact of patient mutations on the structural integrity of *hs*PURA had to rely on a human homology model derived from *dm*PURA (Reijnders *et al*., 2018). Since *Drosophila* and human PURA have a rather moderate sequence identity of 46%, we decided to establish a better experimental basis for predicting the structural effects of patient mutations and determined the crystal structure of the fragment P215-K280 (i.e. *hs*PURA III) at 1.7 Å resolution (**Fig 3A** and **Appendix Table S2**).

**Figure 3.**
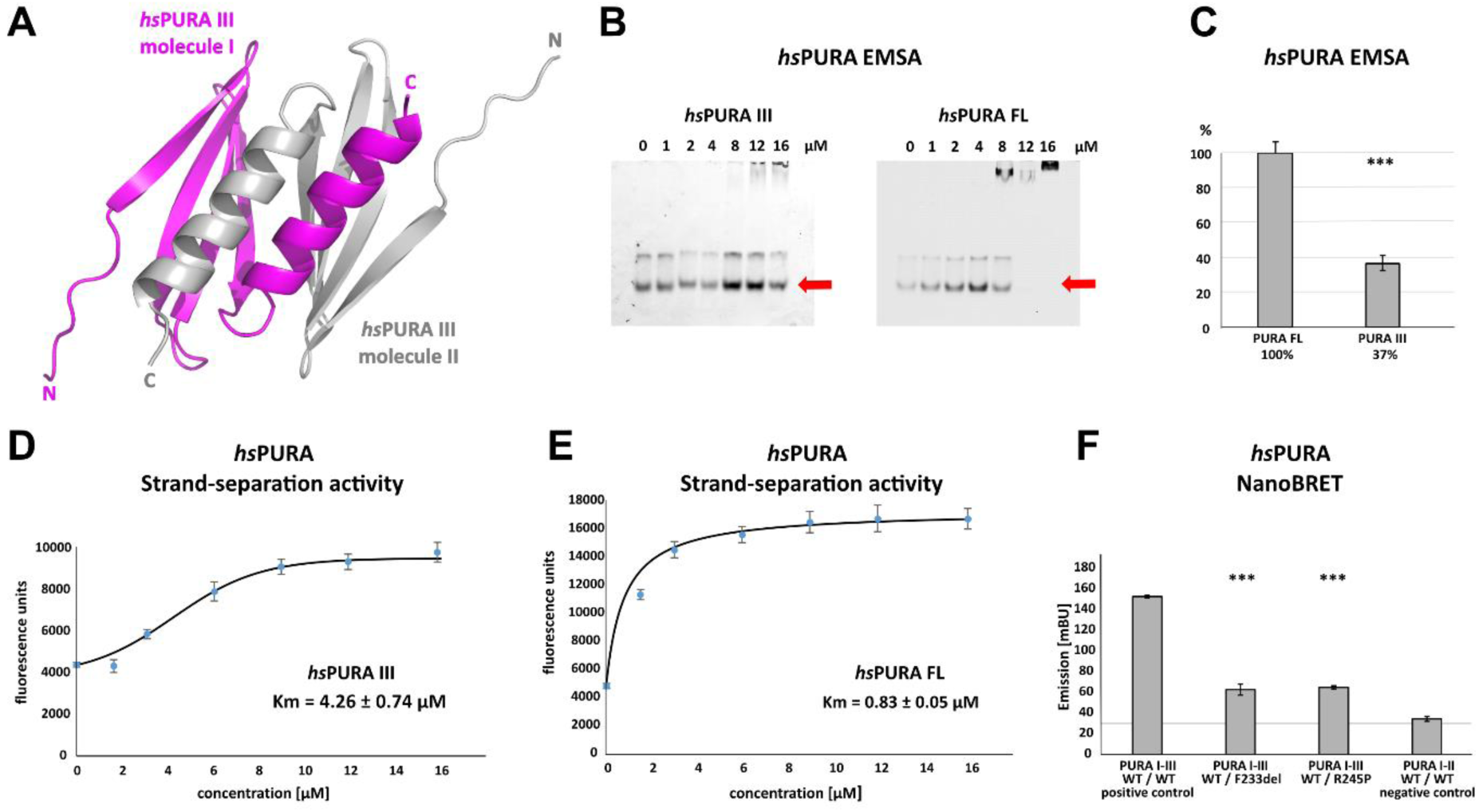
Crystal structure of C-terminal PUR domain from *hs*PURA and effects of mutations on its function. **A** Crystal structure of *hs*PURA III at 1.7 Å resolution (PDB ID: 8CHW). Two *hs*PURA III molecules (magenta and grey, respectively) form a homodimer. Like other PUR domains (e.g. *hs*PURA I-II), *hs*PURA repeat III consists of four β-sheets and one α-helix. **B** In electrophoretic mobility shift *assays (EMSA)* the interaction of *hs*PURA FL and *hs*PURA III with a 24-mer (CGG)_8_ RNA fluorescently labeled with Cy5 fluorophore at 5’end was observed. The amount of the RNA was kept constant at 8 nM while the protein concentration increased from 0 to 16 μM. The unbound RNA used for quantification (see E) is indicated with red arrows. **C** Apparent affinities derived from EMSAs (C) indicate that the C-terminal PUR domain is also able to interact with the nucleic acids. Pairwise t-tests of the *hs*PURA fragments showed significantly lower RNA binding compared to the full-length protein (*hs*PURA III: p = 9.6*E*-4). Three replicates were measured for each experiment and protein variant, and the standard deviations have been calculated and shown as bars. **D**, **E** Strand-separation activity of *hs*PURA. The graphs show the averaged values of three independent experiments as dots with standard deviations as error bars. **D** The *hs*PURA III fragment shows sigmoidal increase of the ssDNA concentration measured as a fluorescent signal. For the quantification of the sigmoidal curves, we utilized Boltzmann function implemented in the Origin software. Calculated x_0_, which corresponds to the Km in this assay yields 4.26 ± 0.74 µM. **E** Strand-separation activity of full-length *hs*PURA was calculated with Michaelis-Menten kinetic, yielding an average Km value of 0.83 ± 0.05 µM, respectively. For both *hs*PURA samples at least three independent measurements have been performed. **F** NanoBRET experiments in HEK293 cells with different *hs*PURA fragment-expressing constructs. Milli-BRET units (mBU) were measured for dimerization of *hs*PURA I-III with *hs*PURA I-III, *hs*PURA I-III F233del, and *hs*PURA I-III R245P. Pairwise t-tests of the mutant *hs*PURA I-III versions F233del and R245P yielded significantly lower signals compared to the wild-type protein (*hs*PURA I-III F233del: p = 3.3*E*-4; *hs*PURA I-III R245P: p = 1.5*E*-7), indicating impaired interactions between the proteins. Black horizontal line shows reference of mBU obtained for *hs*PURA I-II as negative control. Of note, since the BRET signal is the ratio of donor and acceptor signal, it does not require normalization for expression levels. Asterisks in (B) and (G) indicate significance level: *** for p ≤ 0.001.

This fragment corresponds to the domain boundaries suggested by the previously solved invertebrate *dm*PURA III structure (Weber *et al*., 2016). The structure of the C-terminal PUR domain possesses the typical four antiparallel β-strands and one α-helix. Like in the previously published *dm*PURA III structure (Weber *et al*., 2016), the C-terminal PUR domain of *hs*PURA is built from two PUR repeats III of independent protein chains that interact with each other to form an intermolecular homodimer. The *hs*PURA III dimer shows high structural similarity to *dm*PURA III with r.m.s.d. of 1.26 Å for 125 superimposed Cα atoms and 50% of sequence identity (**Appendix Fig S6A**).

### Dimerization domain of *hs*PURA shows nucleic acid binding and unwinding activities

PUR repeats III adopt a classical PC4-like fold, suggesting that it might also bind to nucleic acids. Analysis of the crystal structure of the *hs*PURA III homodimer revealed electrostatic surface potentials favorable for interactions with nucleic acids (**Appendix Fig S6B**). When performing gel electrophoresis mobility shift *assays* (EMSAs) we indeed observed RNA binding (**Fig 3B** and **3C**).

It had been previously reported that PURA can separate double-stranded (ds) nucleic acids in an ATP-independent fashion (Darbinian *et al*, 2001; Weber *et al*., 2016). This so-called unwindase activity has also been reported for several other PC4-like domains from different organisms (Janowski & Niessing, 2020). For *dm*PURA it had already been suggested that dsDNA strands are melted by intercalation of the β-ridge between both strands, followed by binding of both single strands to the cavities formed by the curved β-sheets on both sides of the β-ridge (**Appendix Fig S7**) (Weber *et al*., 2016). Since we had already observed RNA binding by PUR repeats III (**Fig 3B** and **3C)**, we examined its potential unwindase activity by utilizing a previously reported *in vitro* unwinding assay (Darbinian *et al*., 2001; Weber *et al*., 2016) with dsDNA 5’-end labelled with a FAM fluorophore and 3’-end labelled with Dabcyl quencher (**Appendix Fig S8).** In this assay, double-stranded DNA results in low fluorescence signal due to the proximity of the quencher to the fluorophore. Upon strand separation their distance and fluorescence signal increase. We indeed observed strand separation in presence of the *hs*PURA III fragment (**Fig 3D**). Of note, RNA binding as well as dsDNA strand separation by full-length *hs*PURA was much more efficient than by the PUR III domain (**Fig 3D** and **3E**), indicating potential cooperativity of the N- and C-terminal PUR domains in unwinding double-stranded nucleic acids.

### PURA syndrome-causing mutations in the C-terminal PUR domain impair dimerization

Next, we used the *hs*PURA III structure (**Fig 3A** and **Table S2**) to perform an *in silico* analysis of the impact of patient-derived mutations F233del and R245P on the structural integrity of the C-terminal PUR domain. While failure to obtain soluble recombinant PURA with F233del mutation prevented a direct biophysical assessment of its dimerization state *in vitro* (**Appendix Table S1**), this observation already indicated a misfolding of the C-terminal PUR domain upon mutation. Consistent with this observation is also the interpretation from homology modeling based on the previously published *dm*PURA III structure (Reijnders *et al*., 2018). There, the mutation was predicted to impair the structural integrity of the C-terminal *hs*PURA dimerization domain. F233 is part of the β-sheet and interacts with the α-helix of the other PUR repeat III within the PUR domain, contributing to dimerization (**Appendix Fig S9A**). Since in the PURA F233del variant, one residue is deleted, the orientation of all subsequent amino acids in the β-strand adopt the opposite orientation, likely resulting in a fold disruption of the β-sheet and possibly in impaired dimerization. In the case of R245P mutation (**Appendix Fig S9B**), the exchange into proline also likely impairs β-strand folding by introducing a kink, which could also impact dimerization.

To provide direct proof for impaired dimerization upon the introduction of patient-derived F233del and R245P mutations, we utilized the FRET-based NanoBRET assay in HEK293 cells **(Fig 3F**). With this assay, we assessed the ability of *hs*PURA bearing the F233del and R245P patient mutations to dimerize with wild-type *hs*PURA by measuring corresponding milli BRET Units (mBU). Of note, this combination of wild-type and mutant PURA protein is likely to recapitulate the heterozygous situation in patients. As a control for background signal, non-dimerizing *hs*PURA I-II was measured (negative control). As a control for positive interaction, *hs*PURA I-III was used. While for *hs*PURA I-III the signal intensities above the background level clearly indicated dimerization (**Fig 3F**), protein fragments with either the F233del or the R245P mutation showed strongly reduced signal intensities. This observation implies impairment of PURA dimerization in patients bearing either of these heterozygous mutations.

### N-terminal PUR domain shows great flexibility in its domain fold

For a better understand of the structural impact of mutations in the N-terminal PUR domain (*hs*PURA I-II), we determined its structure (fragment E57-E212) by X-ray crystallography (**Fig 4A** and **Appendix Table S2**). The N-terminal PUR domain resembles the previously published PURA structure from *D. melanogaster* (Graebsch *et al*., 2009; Weber *et al*., 2016) (*dm*PURA I-II WT) with an average root mean square deviation (r.m.s.d.) of 1.2 Å (**Appendix Fig S10**). However, in contrast to this invertebrate structure, we observed for *hs*PURA I-II that the four molecules within its asymmetric unit of the crystal lattice showed marked differences in its loop regions and in particular at the edges of the β-sheets (**Fig 4B**). This suggested greater structural flexibility than average domain folds and most likely than its *D. melanogaster* ortholog.

**Figure 4.**
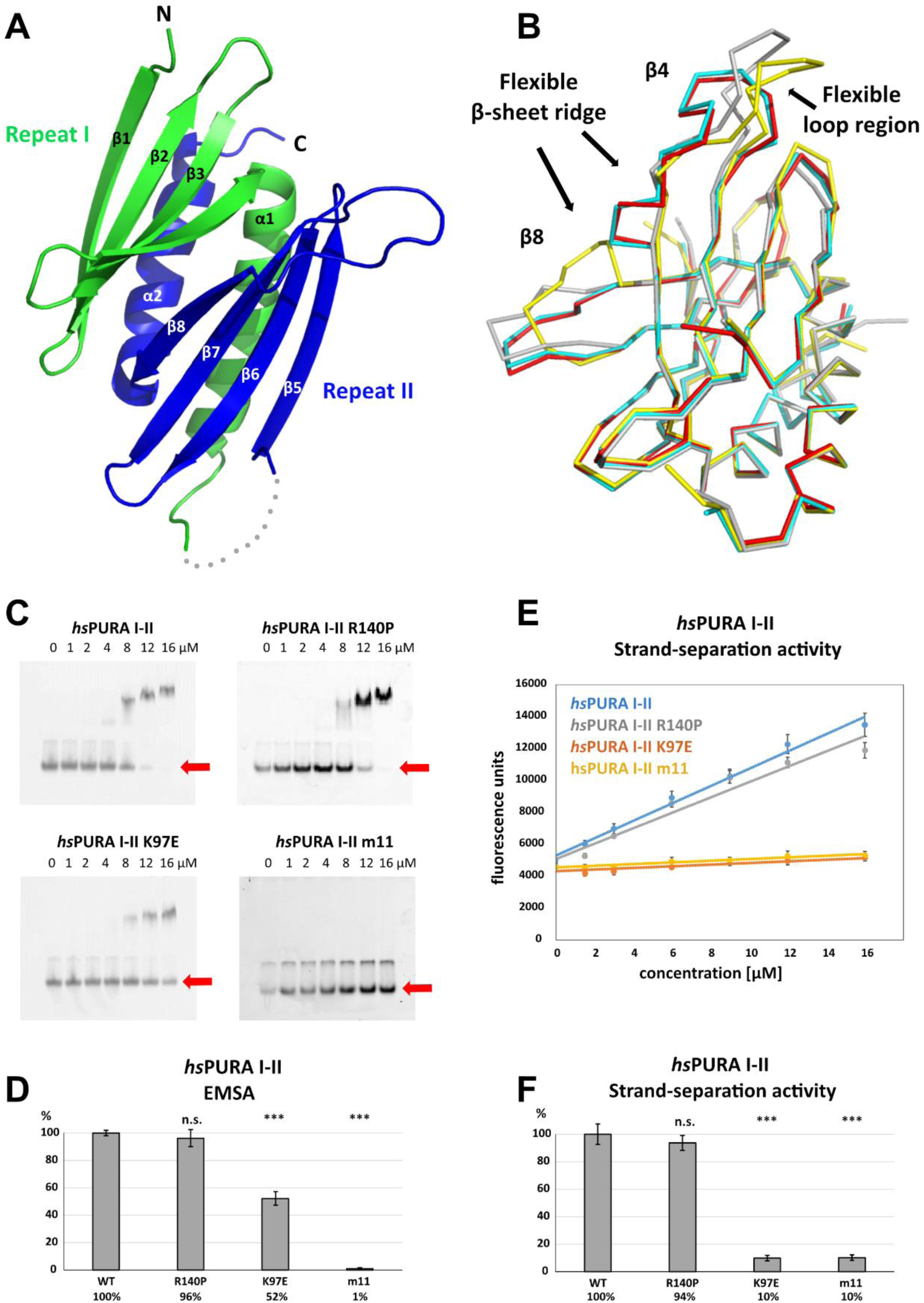
Structure and nucleic-acids interaction of the N-terminal PUR domain. **A** Crystal structure of wild-type *hs*PURA repeat I (green) and II (blue) at 1.95 Å resolution. Protein fragment for which no electron density was visible is shown as a grey dotted line. **B** Ribbon presentation of the overlay of all four chains (shown in different colors) of *hs*PURA I-II in the asymmetric unit of the crystal. There are three regions showing greater differences in folding between these individual molecules, indicating that they are flexible and likely adopt several conformations also in solution. **C** In electrophoretic mobility shift assays, (EMSA) the interaction of different *hs*PURA I-II variants with 24-mer RNA (CGG)_8_ labeled with Cy5-fluorophore was observed. The amount of labeled RNA was kept constant at 8 nM while the protein concentration increased from 0 to 16 μM. The unbound RNA was used for quantification (see D) and is indicated with the red arrow. Above the unbound RNA a second band of free RNA is visible, which might constitute a different conformation or RNA dimer. **D** Quantification of the RNA interactions of different *hs*PURA I-II variants from EMSAs shown in (C). The RNA-binding affinity of wild-type *hs*PURA was normalized to 100%. Pairwise t-test was used to assess differences in RNA binding compared to the *hs*PURA I-II. The mutants K97E and m11 showed significantly lower relative affinities (p = 1.5E-05 and p = 4.6E-09, respectively). In contrast, the mutation R140P did not alter binding affinity (p = 0.5). **E** Strand-separation activity of different *hs*PURA variants. For each *hs*PURA sample at least three measurements have been performed. The graphs show the averaged values as points as well as the standard deviations as error bars. The strand-separation activity shows linear increase within the used concentration range. **F** Quantitative representation of strand-separating activity of the *hs*PURA I-II variants. The strand-separating activity of *hs*PURA I-II was quantified from the slope in (E) and normalized to 100%. Except for R140P (p = 0.07), pairwise t-tests of the *hs*PURA I-II mutants showed significantly lower relative activity (K97E p = 3.9E-16 and m11 p = 9.2E-16) than *hs*PURA I-II. For each experiment and each protein variant three replicates were measured, the standard deviations were calculated and are shown as bars. Asterisks in (D) and (F) indicate significance level: * for p ≤ 0.05, *** for p ≤ 0.001; n.s. for p > 0.05.

### Effects of patient mutations in the N-terminal PUR domain on RNA binding

To assess the interaction of *hs*PURA I-II with nucleic acids and to understand how selected patient-derived mutations affect this function, we performed EMSAs for this protein fragment (**Fig 4C** and **4D**). While *hs*PURA I-II quantitatively bound a 24-mer CGG-repeat RNA, the *hs*PURA I-II K97E mutant protein showed moderately but significantly decreased RNA binding (**Fig 4C** and **4D**). Impaired RNA binding upon K97E mutation is not surprising as a change of the surface electrostatic potential from positive to negative may impact its interaction with nucleic acids (**Appendix Fig S6B**). In contrast, *hs*PURA I-II R140P did not show impaired RNA binding (**Fig 4C** and **4D**). This finding indicates that R140P might be a benign mutation and does not have disease-causing properties. As a negative control, the N-terminal PUR domain containing eleven selected point mutations was employed (*hs*PURA I-II m11; **Appendix Fig S2**). Since these mutations were introduced to abolish the interaction of *hs*PURA with nucleic acids, this mutant protein showed no detectable RNA binding in EMSA.

### PURA-dependent strand separation of dsDNA is affected by disease-causing mutations

We recapitulated the previously reported unwindase activity (Darbinian *et al*., 2001; Weber *et al*., 2016) with human PURA I-II and asked if patient-derived mutations in this domain impair PURA’s strand-separating activity. *In vitro* unwinding experiments with dsDNA were performed for *hs*PURA I-II and compared to the patient mutants *hs*PURA I-II K97E and *hs*PURA I-II R140P as well as the *hs*PURA I-II m11 variant. While *hs*PURA I-II showed considerable strand-separating activity in this assay, the nucleic acid-binding deficient m11 mutant variant expectedly failed to separate dsDNA (**Fig 4E** and **4F**). With an effect considerably stronger than observed for RNA binding (**Fig 4C** and **4D**), the *hs*PURA I-II K97E variant showed a loss of strand-separation activity by 90% (**Fig 4E** and **4F**). In contrast, *hs*PURA I-II R140P mutant showed wild-type-like separation of dsDNA (**Fig 4E** and **4F**), further supporting our previous interpretation from RNA-binding analyses that the mutation R140P is likely functional.

### CD spectroscopic measurements indicate impaired domain folding upon mutations

In order to confirm the structural integrity of the soluble *hs*PURA variants, we performed circular dichroism (CD) spectroscopy experiments. The spectra for both, *hs*PURA I-II as well as *hs*PURA I-II R140P mutant showed a typical alpha-beta profile (**Fig 5A**), suggesting a proper folding of the protein. In contrast, the spectrum for *hs*PURA I-II K97E indicated a strongly altered secondary-structure content. This observation was unexpected as we had predicted that this surface mutation would not affect the protein’s structural integrity (this study and (Reijnders *et al*., 2018). It also indicates that more detailed structural information of *hs*PURA I-II is needed to better understand its susceptibility to disease-causing mutations of surface-exposed residues.

**Figure 5.**
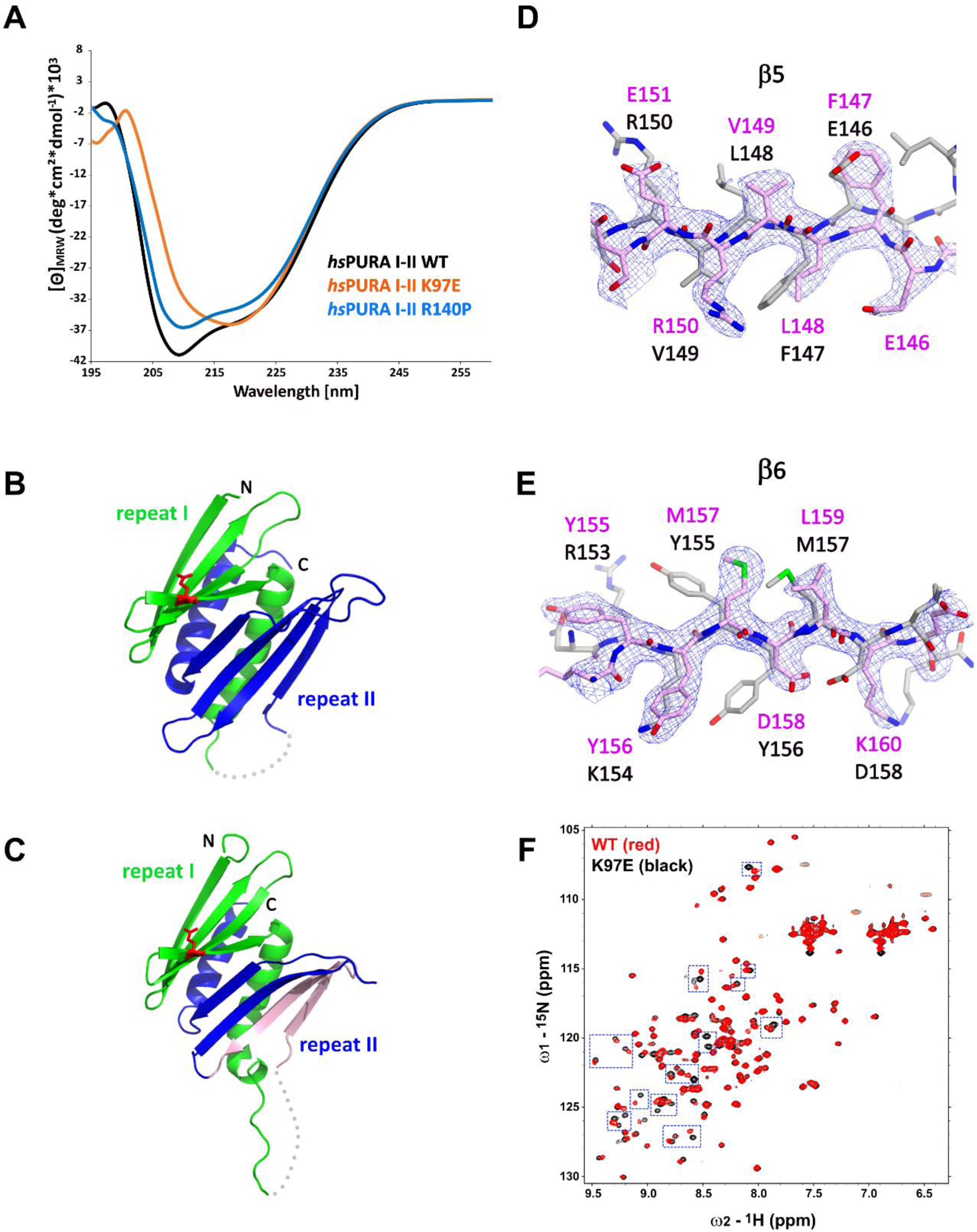
Structural analysis of *hs*PURA I-II bearing the K97E patient mutation. **A** Circular dichroism (CD) spectroscopic analyses of *hs*PURA I-II, K97E, and R140P mutant. Shown are the mean of three CD measurements for each of the proteins (n = 3). The spectra of *hs*PURA I-II K97E show a very different profile, indicating an altered fold of this mutant compared to the wild-type form. **B, C** Crystal structures of *hs*PURA K97E repeat I (green) and II (blue) at 2.45 Å resolution, showing a high overall similarity to *hs*PURA (**Fig 4A**). Two independent *hs*PURA I-II K97E chains (A and B) in the asymmetric unit are shown in (**B**) and (**C**), respectively. The mutated amino acid K97E is indicated as red sticks. **D, E** Positional shifts of amino acids in β5 (**D**) and β6 (**E**) strands of the chain B in the crystal structure of *hs*PURA I-II K97E. 2Fo-Fc electron density map (contour 1σ) is shown for the selected fragment of β5 and β6 strands in chain B (light pink). The superimposed chain A (grey) shows a register shift in chain B by +1 amino acid in the β5 strand and by +2 in the β6 strand. **F** NMR experiment with wild-type *hs*PURA I-II and *hs*PURA I-II K97E. Overlay of the ^1^H, ^15^N-HSQC spectra of *hs*PURA I-II (red) and K97E (black). Blue boxes indicate examples of changes between both spectra.

### *hs*PURA bearing mutation R140P with uncertain clinical significance shows wild-type-like domain folding

The mutation R140P had been previously described to be of uncertain clinical significance and it remained to be shown if this mutation affects the protein’s domain folding. CD spectra had already suggested that this mutation has no dramatic effect on the overall domain folding (**Fig 5A**). To assess if this mutation has a rather local effect on the structural integrity of PURA that may not be detectable by CD spectroscopy, we solved the crystal structure of *hs*PURA I-II R140P at 2.15 Å resolution (**Appendix Fig S11A, S11B** and **Table S2**). The crystals of R140P mutant protein exhibited the same space group, unit cell parameters, and crystal packing as the wild-type protein (**Appendix Table S2**). The R140P mutation is located at the end of the flexible linker between PUR repeat I and II connecting helix α1 to the strand β5 (**Fig 1A**, **Appendix Fig S11A** and **S11B**). Because of its position in a flexible region, the mutated residue is partially visible only in one out of four PURA chains in the asymmetric unit. The overlap of the R140P mutant structure with the PURA I-II WT fragment confirmed our previous observation about the unusually high flexibility of the strands β4 and β8 forming the β-ridge (**Appendix Fig S11A** and **S11B**). Since no major structural differences were visible between *hs*PURA I-II WT and R140P mutant (average r.m.s.d. of 0.89 Å for 133 Cα atoms aligned, **Appendix Fig S11C**), we concluded that this mutation is unlikely to impair the structural integrity of *hs*PURA.

### *hs*PURA I-II with the patient mutation K97E reveals promiscuous domain folding

To understand how exactly the surface-exposed K97E mutation alters the domain folding of PURA (**Fig 5A**), we solved the crystal structure of *hs*PURA I-II K97E (**Fig 5B-E**, **Appendix Table S2)**. The asymmetric unit of the crystal contained two molecules of *hs*PURA I-II K97E and showed good overall fold similarity to the *hs*PURA I-II structure (**Fig 5B** and **5C**). However, an overlay of both molecules of the K97E-mutant protein from the asymmetric unit also revealed unusual differences between them with r.m.s.d of 1.48 Å for 125 superimposed Cα atoms. While one of these molecules (chain A, **Fig 5B**) showed a fold very similar to *hs*PURA I-II (average r.m.s.d. 1.09 Å, **Appendix Fig S12A**), the second molecule (chain B) displayed much greater structural changes (average r.m.s.d. 1.61 Å, **Fig 5C** and **Appendix Fig S12A**). These differences were especially pronounced in PUR repeat II (**Fig 5D-E**), where we observed a shift of register in the β-sheet by one amino acid in strand β5 (**Fig 5D**) and by two amino acids in strand β6 (**Fig 5E**). More C-terminally, the loop linking β6 and β7 is shorter by two residues, thereby correcting again the sequence register within the next β-strands (β7 and β8).

Already the different molecules of wild-type *hs*PURA I-II in the crystal unit cell revealed considerable conformational freedom (**Fig 4B**, **Appendix Fig S12B** and **S12C**), not only for narrow movements of the loops but also in the folding of the β-strands. For instance, in some of the molecules in the asymmetric unit of the wild-type *hs*PURA I-II crystal, β4 and β8 strands formed bulges (**Fig 4B**, **Appendix Fig S12B** and **S12C**). From all the areas of the structure, the β-ridge appears to be structurally the most divergent part. It is surprising that the K97E point mutation, which is located on the β4 strand of the β-ridge, induces the observed unusual change in the β-sheet arrangement (**Fig 5C-E**) without destroying the domain’s overall scaffold. Together these findings indicate great structural flexibility of the N-terminal PUR domain and promiscuity in folding, which results in misfolding upon mutation of a surface-exposed residue. While CD measurements suggest a difference in folding between wild-type and K97E mutated PURA (**Fig 5A**) and the crystal structure of *hs*PURA I-II K97E indicates folding promiscuity (**Fig 5C-E**), these data do not fully exclude unfolding of the domain in solution. To distinguish between a mutation-induced unfolding and an altered folded state, we performed NMR measurements with the ^15^N-isotope-labeled samples of wild-type and K97E mutant *hs*PURA I-II (E57-E212 fragment). First, ^1^H,^15^N-correlation spectra (HSQC) of both wild-type and K97E constructs showed a clear presence of the structured core, as evidenced by the well-dispersed resonances, with a significant number of residues adopting strand conformation (^1^H 8.5 - 9.5 ppm) (**Fig 5F**). Next, we carefully compared the two HSQC spectra for individual resonances/residues. When the two spectra were overlaid, we observed that a significant number of resonances, especially in the region corresponding to the β-strand, were perturbed, which is clearly more extensive than just affecting the residues adjacent to the mutational site of residue 97 (**Fig 5F**). Finally, we performed the ^15^N-heteronuclear NOE experiment on the K97E mutant to measure the residue-level backbone dynamics (**Appendix Fig S13A**). The overlaid spectra of the absence (control) and presence (NOE) of ^1^H-saturation shows most resonances retaining their signal intensities (rigid, hetNOE ∼ 0.8), while the resonances corresponding to the dynamic regions, likely N-/C-terminal & loop regions, with significantly reduced signal intensities or even negative (flexible). Importantly, all the resonances perturbed upon K97E mutation remained in the presence of ^1^H-saturation, confirming that the structural rearrangement does not induce protein unfolding. Together, this shows that the K97E mutation does not induce global unfolding of *hs*PURA I-II, but rather triggers structural rearrangements mostly in the β-strand region.

To obtain a better understanding of the intrinsic dynamics of *hs*PURA I-II we performed *in silico* molecular dynamics simulations. The *hs*PURA I-II WT and K97E showed a similar overall profile (**Appendix Fig S13B**), where most of the flexibility or fluctuations are concentrated in the loops connecting strands β3 with β4 and β7 with β8. However, there are some statistically significant differences between the two proteins. The most significant differences concentrate on the two ends of the α1 helix, as well as the β5, β6, and β7 segments. In these regions, the K97E mutant shows higher flexibility than the WT, with the N-terminal part of the α1 helix and the C-terminal end of β5 and β6 having the greatest gap between the two variants. The movie (S1 for *hs*PURA I-II WT and S2 for *hs*PURA I-II K97E) of the simulation confirms a large degree of fluctuation in this area, which can adopt many different conformations in both WT and mutant. Finally, there is one region where the WT displays a higher flexibility than the mutant, the loop connecting β7 with β8 (residues 175 to 185), although only a few of those differences are significant. Of note, while these simulations are consistent with the increased flexibility suggested by the crystal structure, the time window of these simulations was too short to observe alterations between the two folding states as seen in this experimental structure.

## DISCUSSION

Currently, more than 270 different pathogenic mutations in the *PURA* gene have been identified in over 600 patients with confirmed PURA syndrome (personal communication: PURA Foundation Australia). The aim of this study was to understand how mutations causing this neurodevelopmental disorder affect the functional and structural integrity of the *hs*PURA protein. Towards this goal, we performed cellular, biophysical, and structural analysis with wild-type *hs*PURA as well as with versions of *hs*PURA bearing representative patient-derived mutations.

To understand how PUR-syndrome-causing mutations affect the subcellular localization of *hs*PURA, we performed immunofluorescence microscopic analyses in HeLa cells by overexpressing full-length *hs*PURA as well as variants bearing a disease-causing mutation in each of the three PUR repeats, i.e. K97E, I206F, and F233del. Consistent with previous reports (Daigle *et al*., 2016; Markmiller *et al*., 2018), we observed that *hs*PURA co-localizes to stress granules in HeLa cells (**Fig 1B** and **1C**). As we did not observe significant changes in the association of patient-related mutations of hsPURA to stress granules, it is suggested that this feature may not constitute a major hallmark of the PURA syndrome. It should be noted however that this interpretation must be considered with some caution as the experiments were performed in a PURA wild-type background.

What is more, the knockdown of PURA resulted in reduced sizes of those stress granules (**Appendix Fig S4**), indicating that PURA indeed does have an influence on these granules. Future experiments will be necessary to unambiguously clarify the importance of PURA’s localization to stress granules for the etiology of PURA syndrome.

Very recently, *hs*PURA was reported to localize to P-bodies and that a PURA knockdown leads to a significant reduction in HeLa cells (Molitor *et al*., 2023). While we recapitulated co-localization of *hs*PURA to P-bodies (**Fig 2**), *hs*PURA K97E and F233del mutants failed to fully localize to P-bodies. Since the F233del mutation impairs the dimerization of *hs*PURA (**Fig 3F**), this finding suggests the requirement of dimerization for efficient P-body association.

To address the question of what drives *hs*PURA accumulation in these phase-separated granules, we utilized a *hs*PURA version lacking RNA-binding activity (m11, **Appendix Fig S2**). This mutant failed to localize to stress granules or P-bodies, indicating that RNA binding is essential for both co-localization events. A potential RNA-dependent mechanism to recruit *hs*PURA into stress granules or P-bodies could be liquid phase separation of *hs*PURA when bound to RNAs. *In vitro* experiments testing this potential property of *hs*PURA failed to yield phase-separated droplets. In summary, these findings indicate that RNA binding of *hs*PURA is important for its co-localization into stress granules and P-bodies, while we have no evidence that phase separation of *hs*PURA plays a role.

In a recent study, it was shown that a *hs*PURA knock-down in HeLa and in fibroblast cells resulted in down-regulation of the essential P-body proteins LSM14a and DDX6 and, most likely as a direct consequence, in strongly impaired P-body formation (Molitor *et al*., 2023). In light of this report, our finding that patient-derived mutations impair the P-body association of *hs*PURA (**Fig 2**) provide further evidence for a potential role of P-body localization of *hs*PURA for the etiology of PURA syndrome.

It has been reported that protein domains with PC4-like fold interact with RNA and DNA and are responsible for the separation of the double-stranded nucleic acids (Darbinian *et al*., 2001; Weber *et al*., 2016) (reviewed in (Janowski & Niessing, 2020). In this study, we show that both, the N- and C-terminal PUR domains of human PURA can separate double-stranded nucleic acids (**Fig 4E**, **4F** and **3D**) and to bind single-stranded nucleic acids (**Fig 4C**, **4D, 3B** and **3C**). We also tested patient-derived mutant versions of *hs*PURA to find out whether they exhibit impaired interaction with nucleic acids. For the K97E mutated version, we observed in EMSAs a reduced affinity to RNA (**Fig 4C** and **4D**). Similar behavior of the K97E mutant was observed in the strand-separation assay where it showed an almost complete loss of activity (**Fig 4E** and **4F**). The latter is of interest since previous experiments had demonstrated that strand separation contributes to the neuroprotective activity of PURA in a *Drosophila* FXTAS disease model (Weber *et al*., 2016). It is therefore also probable that loss of strand-separation activity contributes to cellular abnormalities in *hs*PURA patients harboring the K97E mutation.

To establish a structural framework for these observations, we solved the crystal structures of *hs*PURA III (**Fig 3A**) and of *hs*PURA I-II (**Fig 4A**). These human structures were overall very similar to previously published crystal structures from *dm*PURA (**Appendix Fig S6A** and **S10**) (Graebsch *et al*., 2009; Weber *et al*., 2016). Based on the crystal structures of *hs*PURA III (**Fig 3A**) we were able to predict folding defects in the C-terminal PUR domain (*hs*PUR III) that are caused by the patient-derived mutations F233del and R245P (**Appendix Fig S9**). Deletion of phenylalanine 233 within the β2-strand changes the orientation of all subsequent side chains within the β-sheet and likely destroys the interaction network required for the assembly of this secondary structure. A mutation of arginine 245 into proline likely disrupts β-strand folding by introducing a kink. Since repeat III mediates homodimer formation through swapping of its α-helices, the dimerization of *hs*PURA is potentially affected by folding defects induced by either the F233del or the R245P mutation. Indeed, we could experimentally confirm by nanoBRET measurements in HEK293 cells that dimerization of *hs*PURA I-III F233del and *hs*PURA I-III R245 is impaired when compared to wild-type *hs*PURA (**Fig 3F**).

Determination of the crystal structure of *hs*PURA I-II revealed larger flexible regions than the previously reported invertebrate structures (Graebsch *et al*., 2010; Graebsch *et al*., 2009; Weber *et al*., 2016), with considerable differences in the four molecules of *hs*PURA I-II in the asymmetric unit of the crystals (**Fig 4B**, **Appendix Fig S12B** and **S12C)**. These observations provide clear evidence that in particular strands β4 and β8 of the N-terminal PUR domain are dynamic and can adopt considerable local differences within the β-sheet. This flexibility already suggests structural promiscuity of PUR domains. It also indicates that they can at least partially compensate for structural perturbance by mutations, resulting in folded domains of mutant proteins. However, since these mutations were reported to cause PURA syndrome, such modest structural alterations must nevertheless result in functional impairment.

The interpretation of structural flexibility and folding promiscuity of PUR domains found experimental support from the crystal structure of its K97E variant, whose mutated residue has a solvent-exposed side chain (**Fig 5B-E**). The corresponding structure shows in the β5 strand a shift by one (**Fig 5D**) and in the β6 strand by two amino acids (**Fig 5E**) in one of the molecules of the asymmetric unit. Surprisingly, these structural distortions did not result in major unfolding events of this PUR domain. Furthermore, NMR measurements with the ^15^N- and ^1^H-labeled N-terminal PUR domain yielded many spectral differences between the wild-type and K97E variants in particular within their β-sheets (**Fig 5F** and **Appendix Fig S13A**). Despite these considerable structural changes, the NMR data also confirmed that the K97E-mutant protein remains folded in solution and indicates that PUR domains may be able to partially compensate for mutations by folding promiscuity. These findings were further supported by molecular dynamics simulations, indicating high intrinsic motions of the N-terminal PUR domain (**Appendix Fig S13B**).

Instead of hot-spot regions for disease-causing mutations, as reported for other genetic disorders such as neurodevelopmental Rett syndrome (Lombardi *et al*., 2015), mutations of PURA-syndrome patients are found along the entire protein sequence (Johannesen *et al*., 2021; Reijnders *et al*., 2018). Also, in contrast to disorders such as the Rett syndrome, most of these mutations cause the full spectrum of disease symptoms. Our findings suggest that the flexibility and promiscuity of *hs*PURA domain folding render this protein particularly susceptible to mutations, even when amino-acid substitutions impose only moderate changes to the local charge and stereochemistry. This property of PUR domains might offer a structural explanation for why so many mutations across the sequence of *hs*PURA result in the full disease spectrum.

Not in all cases is the pathogenic effect of a given mutation as clear as in the case of K97E. For instance, the replacement of arginine 140 by proline in *hs*PURA was initially reported to cause PURA syndrome (Lee *et al*., 2018). Even though the corresponding patient did show symptoms, the severity and type of symptoms raised doubts about this initial diagnosis. In our assessment, neither the functional *in vitro* experiments nor the structural assessment of *hs*PURA I-II R140P yielded any significant differences from the wild-type protein (**Fig 4C-F** and **Appendix Fig S11**). These experimental findings are consistent with R140P being rather a benign mutation. Furthermore, after the initial description of R140P to be causative for PURA Syndrome (Lee *et al*., 2018), subsequent genomic assessment of a relative of this patient uncovered that this person had the same mutation without any clear symptoms. Consequently, the R140P mutation is now listed in the NCBI ClinVar database to be of “uncertain significance” (https://www.ncbi.nlm.nih.gov/clinvar/variation/192343/). Our data clearly suggest that this mutation is rather benign. In summary, an assessment of the different mutations in *hs*PURA indicates a clear correlation between defects observed in *in vitro* analyses and the reported pathogenicity in patients. Hence, in future such *in vitro* analyses may serve as a diagnostic tool to distinguish pathogenic from benign mutations.

Recently, two publications resorted to *in silico* predicted, AI-based structures to model the potential impact of PURA-syndrome mutations on the structural integrity of *hs*PURA. Instead of assessing the physiologically correct dimer consisting of two PUR repeats, both publications analyzed *in silico*-predicted *hs*PURA half-structure consisting of a single PUR repeat III with incomplete dimerization domain (Dai *et al*, 2023; Lopez-Rivera *et al*, 2022). Such half PURA structures are very unlikely to exist in solution and hence predictions based on them are of very limited value or worse, prone to yield wrong interpretations. We expect that our experimentally determined structures of the human N-and C-terminal PUR domains offer more physiological structural reference points for future assessments of novel *hs*PURA mutations and their potential pathogenicity in patients.

In summary, this study provides the first functional and experimental structural assessment of the effects of PURA-syndrome causing mutations on folding, nucleic-acid binding, and unwinding, as well as on subcellular localization to stress granules and P-bodies. Some of the selected patient variants of *hs*PURA were previously predicted to have folding defects (Reijnders *et al*., 2018). Of note, for all these cases expression tests with recombinant proteins resulted in insoluble samples, indicating that these mutant proteins indeed have impaired folding (**Appendix Table S1**). Whereas predictions of such misfolding events appear reliable, the examples of R140P (benign) and K97E (pathogenic) indicate that structural predictions of surface-exposed residues seem more complicated. In fact, the observed great promiscuity of folding of PUR domains indicates that also moderate surface mutations such as K97E can result in promiscuous folding and the full spectrum of disease symptoms. Our study may serve as proof-of-principle that *in vitro* and structural studies are suitable tools to classify mutations in the *hs*PURA gene as potentially pathogenic or benign.

## MATERIALS AND METHODS

In-depth descriptions of the sample preparation techniques and comprehensive details of the experimental procedures can be found in the Appendix section of this document.

## ACKNOWLEDGMENTS

We acknowledge the use of the X-ray Crystallography Platform at Helmholtz Munich, the Core Facility Bioimaging of the BMC of the Ludwig-Maximilians University Munich, as well as the Bavarian NMR Centre (BNMRZ). We would like to thank Vera Roman for her excellent technical support as well as Melinda Anderson from the PURA Foundation Australia and the PURA Syndrome Foundation for their precious support.

## Author contributions

MP performed structural and biophysical analysis as well as most cell-culture studies, RJ performed X-ray measurements, structural analysis, *in vitro* RNA and DNA binding tests. SB performed staining and analysis of stress granules in HeLa cells under PURA knock-down conditions. SH participated in biophysical analysis as well as cell-based studies. H-S K performed NMR experiments and analysis. TK and AC performed NanoBRET experiments and LM, SB, and JT produced the cell lines. A V-R, R F, E D, and G D F performed and analyzed molecular dynamics simulations. TM performed phase separation experiments. DD, MS and DN supervised and discussed the project. The manuscript was written by MP, RJ, SB, and DN. All authors participated in the discussion, interpreted the results, and commented on the manuscript.

## Data and materials availability

All data are available in the main text or the supplementary materials. Structural models and diffraction data are available at the Protein Data Base (PDB; https://www.rcsb.org/) under the accession codes 8CHT (*hs*PURA I-II), 8CHU (*hs*PURA I-II K97E), 8CHV (*hs*PURA I-II R140P) and 8CHW (*hs*PURA III).

