## Supplemental Figures, Tables, and Methods for "PURA Syndrome-causing mutations impair PUR-domain integrity and affect P-body association"

**Appendix Contents**

Appendix Figure S1-S13
Appendix Table S1 and S2
Appendix Movie S1 and S2
Material and Methods

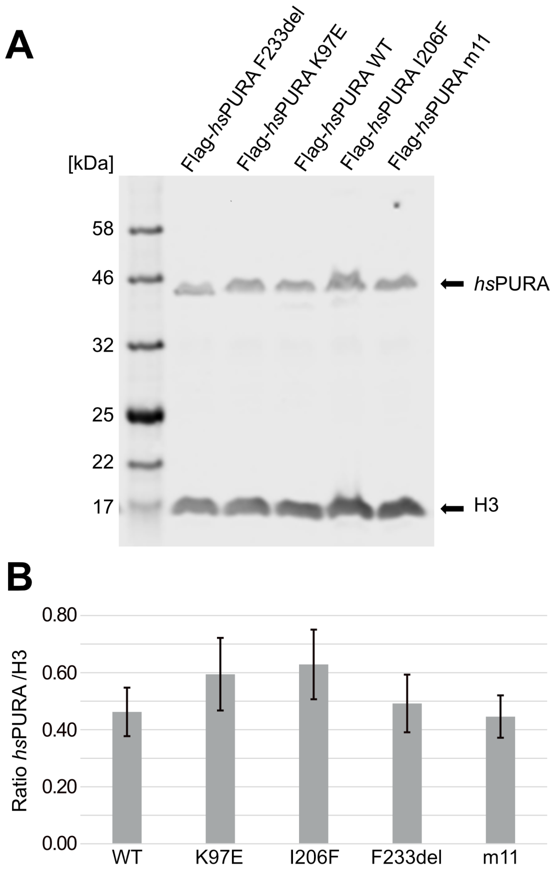


C

**p-values**

| | F233del | I206F | K97E | m11 |
|---|---|---|---|---|
| I206F | 0.29 | - | - | - |
| K97E | 0.49 | 0.80 | - | - |
| m11 | 0.63 | 0.14 | 0.23 | - |
| WT | 0.77 | 0.19 | 0.29 | 0.85 |

**Appendix Figure S1. Quantitative western blot of cell lysate of cell lines used for the immunofluorescence staining assays**.

**A** Used cell lines were doxycycline-induced HeLa overexpression cell lines. Cells were induced 16 h before harvesting, washed twice with PBS and lysed in 2 x SDS Laemmli buffer. For the western blot, commercial antibodies for H3 and Flag were used. H3 was used as a housekeeping marker to quantify the amount of overexpressed protein. The blot shown here is one representative of three replicates (n = 3).

**B** Quantification of the western blots of the IF cell lines. Values represent means ± SD (n=3). No statistically significant difference between cell lines was observed with $p < 0.05$ by ANOVA.

**C** Pairwise t-test confirmed no significant differences between cell lines. All p-values are above 0.05.

**A** ***hs*PURA m11**

**K71A, N80A, K82G, F85A, K87A, K97A, R153A, N162A, R164G, F167A, R169A**

MADRDSGSEQ GGAALGSGGS LGHPGSGSGS GGGGGGGGGG GGSGGGGGGA
PGGLQHETQE LASKRVDIQN **K**RFYLDVKQ**N** A**K**GR**F**L**K**IAE VGAGGN**K**SRL
TLSMSVAVEF RDYLGDFIEH YAQLGPSQPP DLAQAQDEPR RALKSEFLVR
EN**R**KYYMDLK E**N**Q**R**GR**F**L**R**I RQTVNRGPGL GSTQGQTIAL PAQGLIEFRD
ALAKLIDDYG VEEEPAELPE GTSLTVDNKR FFFDVGSNKY GVFMRVSEVK
PTYRNSITVP YKVWAKFGHT FCKYSEEMKK IQEKQREKRA ACEQLHQQQQ
QQQEETAAAT LLLQGEEEGE ED

**B**

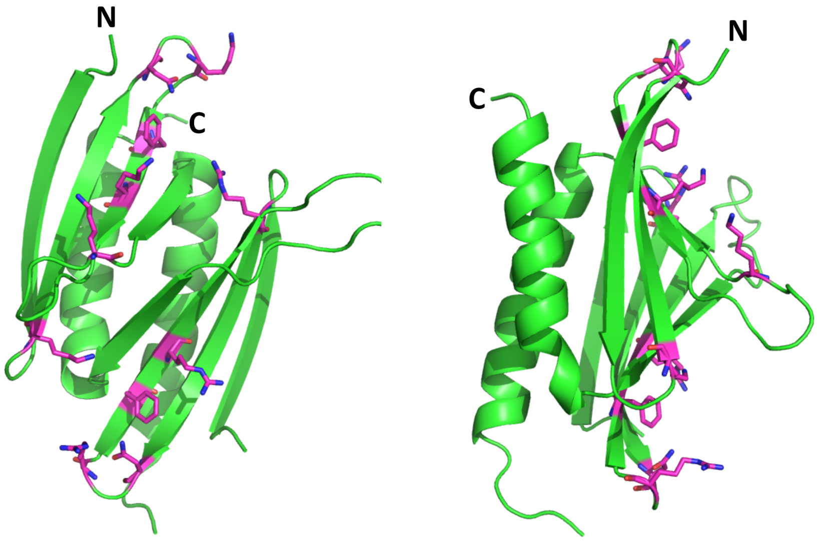


**Appendix Figure S2. RNA-binding-deficient *hs*PURA I-II m11 variant**.

**A** The amino acid sequence of the full-length *hs*PURA protein with the residues constituting the repeats I and II shown in green. Eleven residues that underwent structure-guided point mutations are marked in magenta.

**B** The crystal structure of wild-type *hs*PURA I-II (green ribbon) is shown in two different orientations. The residues which are mutated in the m11 variant are shown as sticks in magenta. These mutations were designed to abolish the interaction of *hs*PURA with nucleic acids.

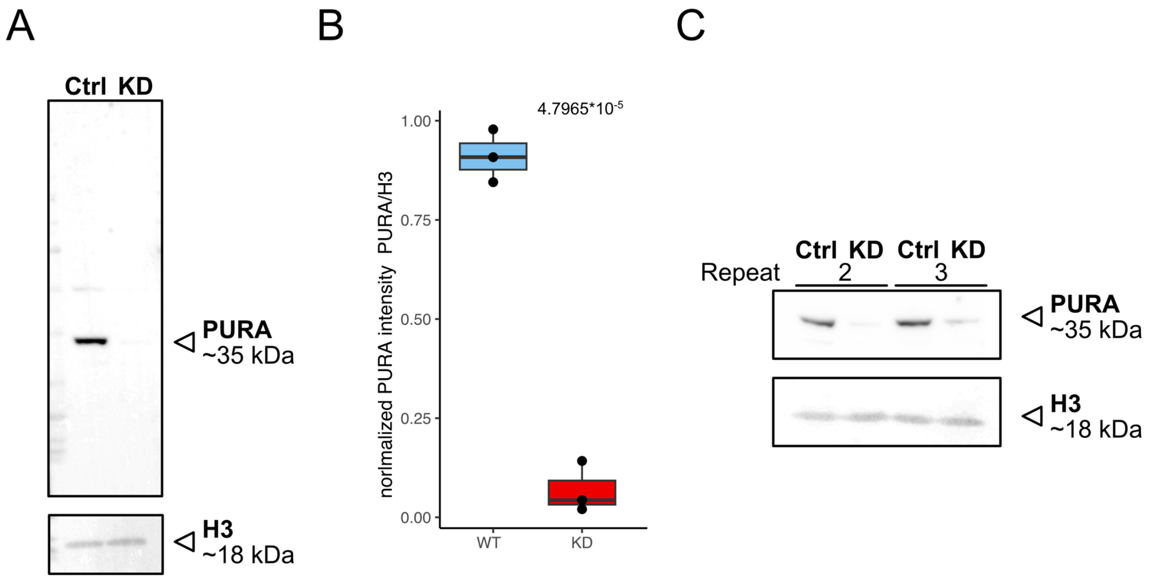


**Appendix Figure S3. Validation of knock down efficiency.**

**A** Western Blot of control (scrambled sirRNA) and PURA knock down HeLa cells anti-PURA and H3 as housekeeping gene.

**B** Quantification of relative PURA intensity normalized to H3 intensity (n=3). Pairwise t-test shows significant differences between cell lines.

**C** Western Blot of Repeat 2 and 3 of KD experiment used for Quantification.

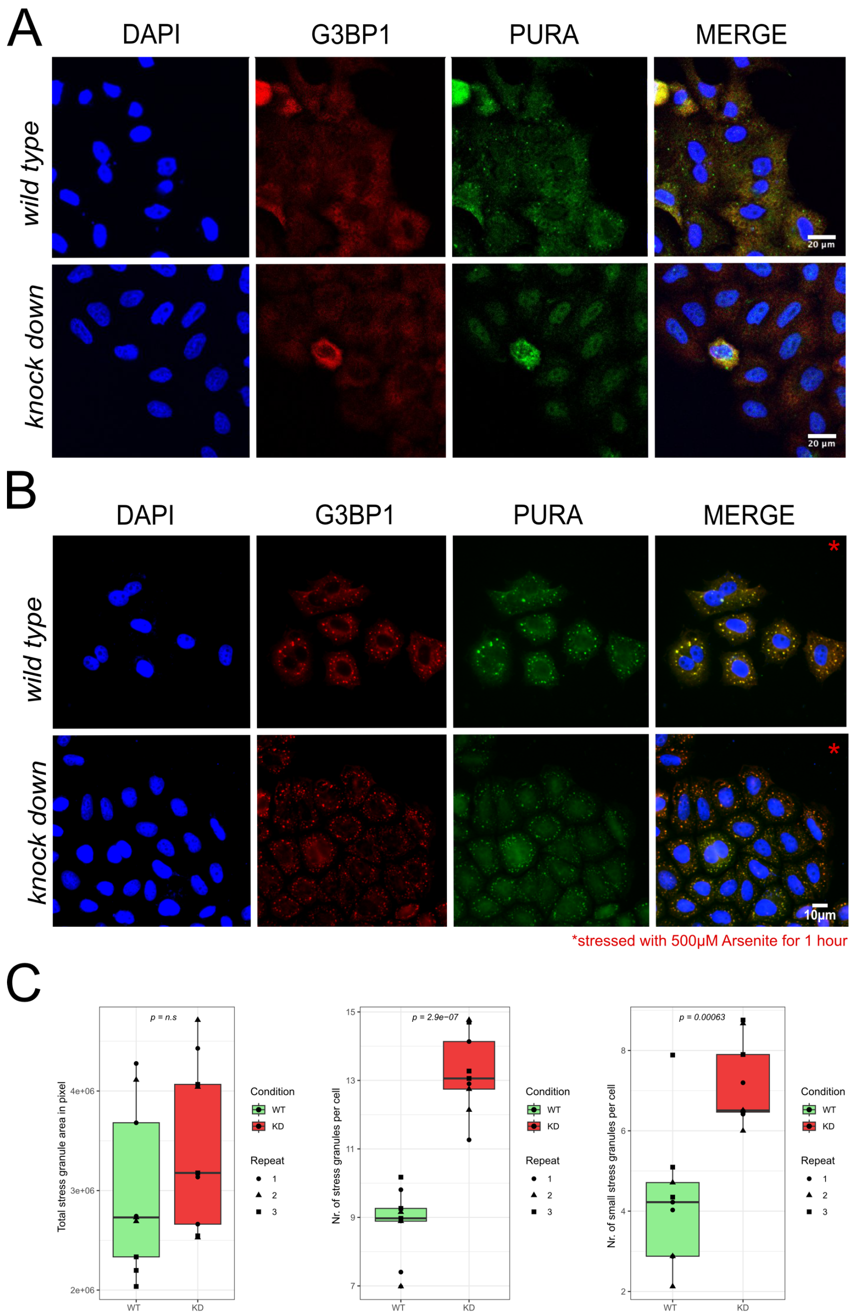


**Appendix Figure S4. Stress-granule formation in HeLa wild-type versus PURA knockdown cells**.

**A** Unstressed HeLa wild-type versus PURA knockdown cells stained with the stress granule marker G3BP1 (red) and PURA (green). No stress granules can be detected by G3BP1 staining.

**B** Arsenite-stressed HeLa wild-type versus PURA knockdown cells stained with the stress granule marker G3BP1 (red) and PURA (green).

**C** Quantification of stress granules per cell in wild-type versus knockdown (KD) HeLa cells under stress conditions shown in (B). While no significant difference can be observed in the overall area of all microscopically visible stress granules (left), significantly more granule number appeared in knockdown cells (middle). Stress granules were defined as entities with ≥ 0.7 µm in diameter and small stress granules as entities with 0.3 µm-0.7 µm in diameter.

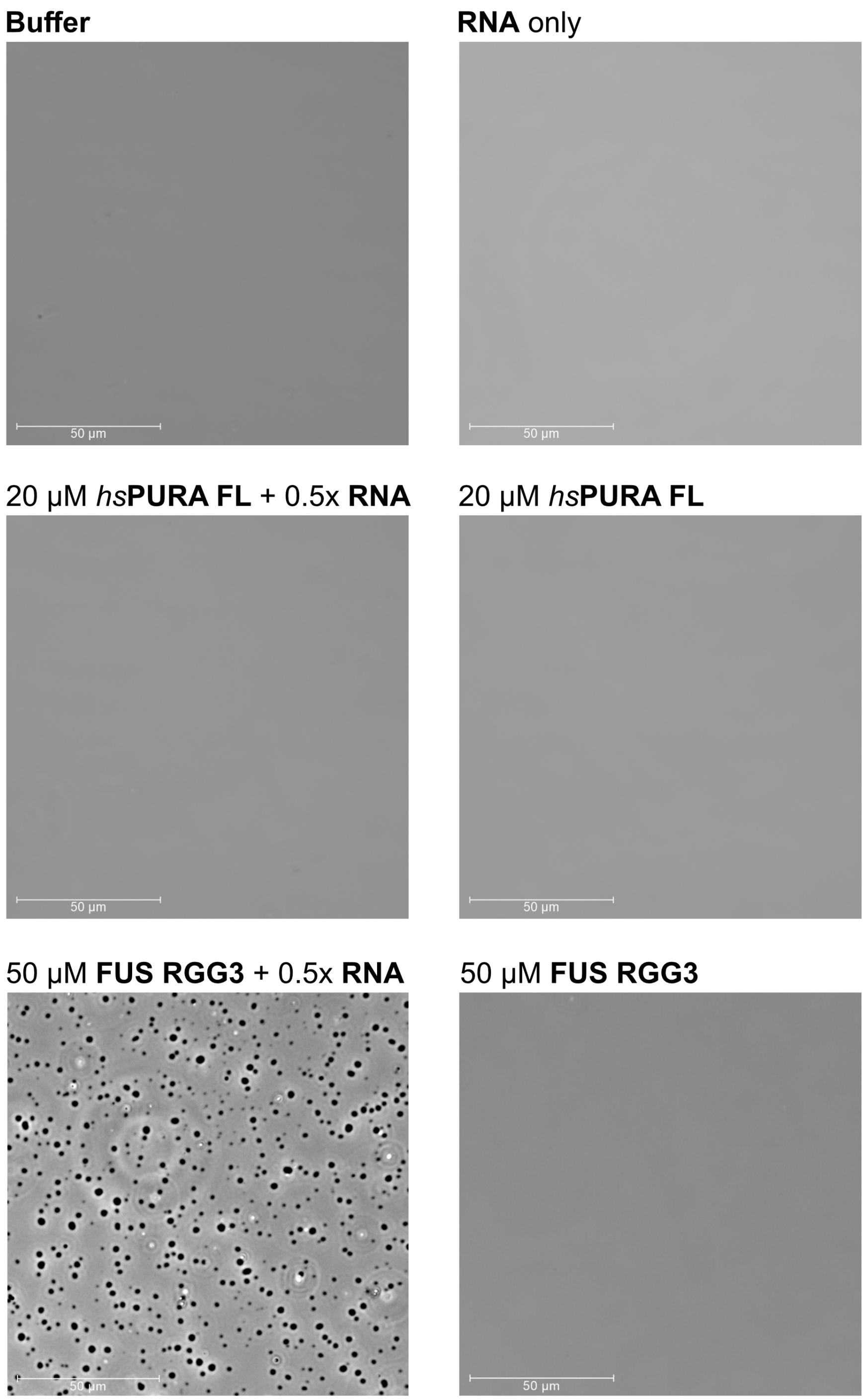


**Appendix Figure S5. *hs*PURA does not undergo phase separation.**

The phase separation experiment was performed with full-lenth *hs*PURA alone and with total HeLa RNA. As a positive control for RNA-dependent phase-separation the unstructured FUS RGG3 protein fragment was used (Dormann *et al*, 2012). No liquid phase separation was observed with *hs*PURA either in absence or presence of RNA.

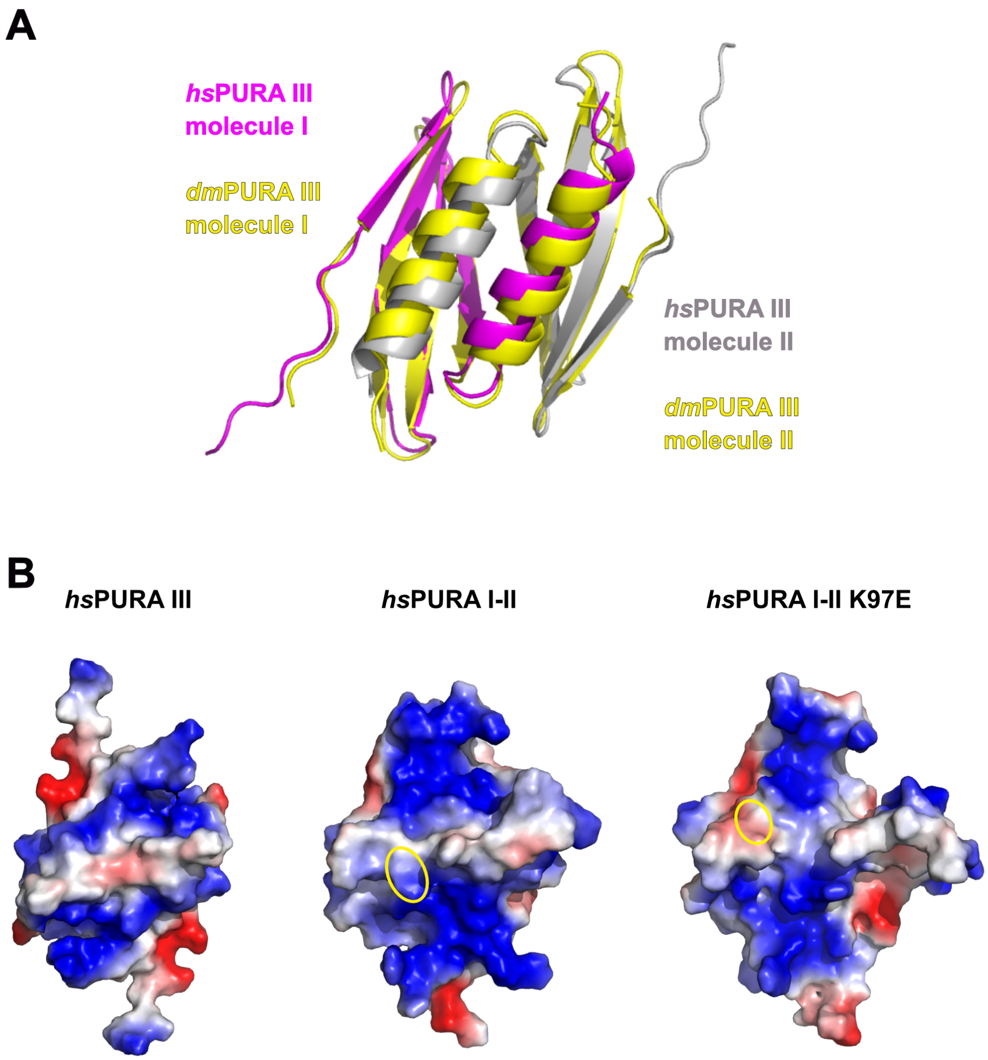


**Appendix Figure S6.**

**A** Superposition of the crystal structure of the *hs*PURA repeat III homodimer at 1.7 Å resolution (PDB ID: 8CHW; chains shown as magenta and gray ribbons) with its *D. melanogaster* homologue *dm*PURA III (PDB ID: 5FGO; both chains shown as yellow ribbons). The *hs*PURA III dimer shows high structural similarity to *dm*PURA III with r.m.s.d. of 1.26 Å for 125 superimposed Cα atoms and 50% of sequence identity.

**B** The surface electrostatic potentials for *hs*PURA III (left), *hs*PURA I-II (middle), and *hs*PURA I-II K97E (right). All three structures are oriented with the nucleic binding sites facing toward the reader. The position of K97 in wild-type *hs*PURA I-II and E97 in *hs*PURA I-II K97E are encircled.

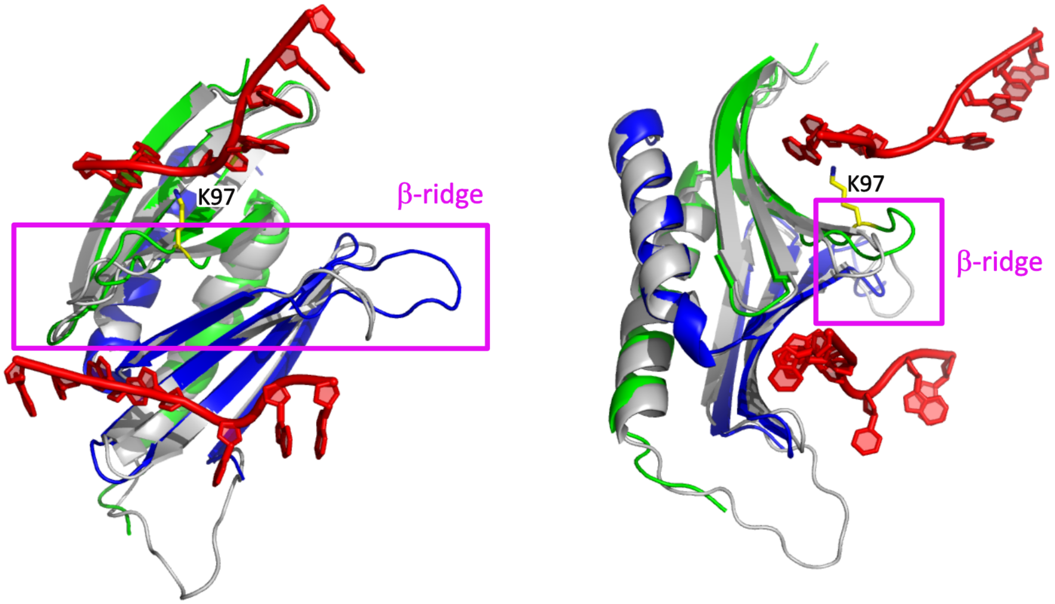


**Appendix Figure S7. Structural features likely contributing to the PURA-dependent unwinding of dsRNA.**

Superposition of the *hs*PURA I-II crystal structure (PDB ID: 8CHT; repeat I in green and II in blue) with the previously published structure of the PURA I-II fragment from *D. melanogaster* (grey ribbon model) in complex with DNA (shown in red) (PDB ID: 5FGP). The structure is presented in two different orientations. The ▫-ridge is highlighted by the magenta square and label. The position of the residue 97 is shown in yellow and labeled accordingly.

FAM-CCAGGGCACTTAAAAAAATTCGCCTGG-Dabcyl

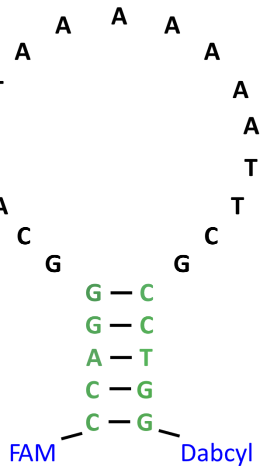


**Appendix Figure S8. The strand-separation activity assay DNA probe.**

The primary and secondary structure of the DNA oligo with 5'-labeled FAM fluorophore and 3'-labeled Dabcyl quencher.

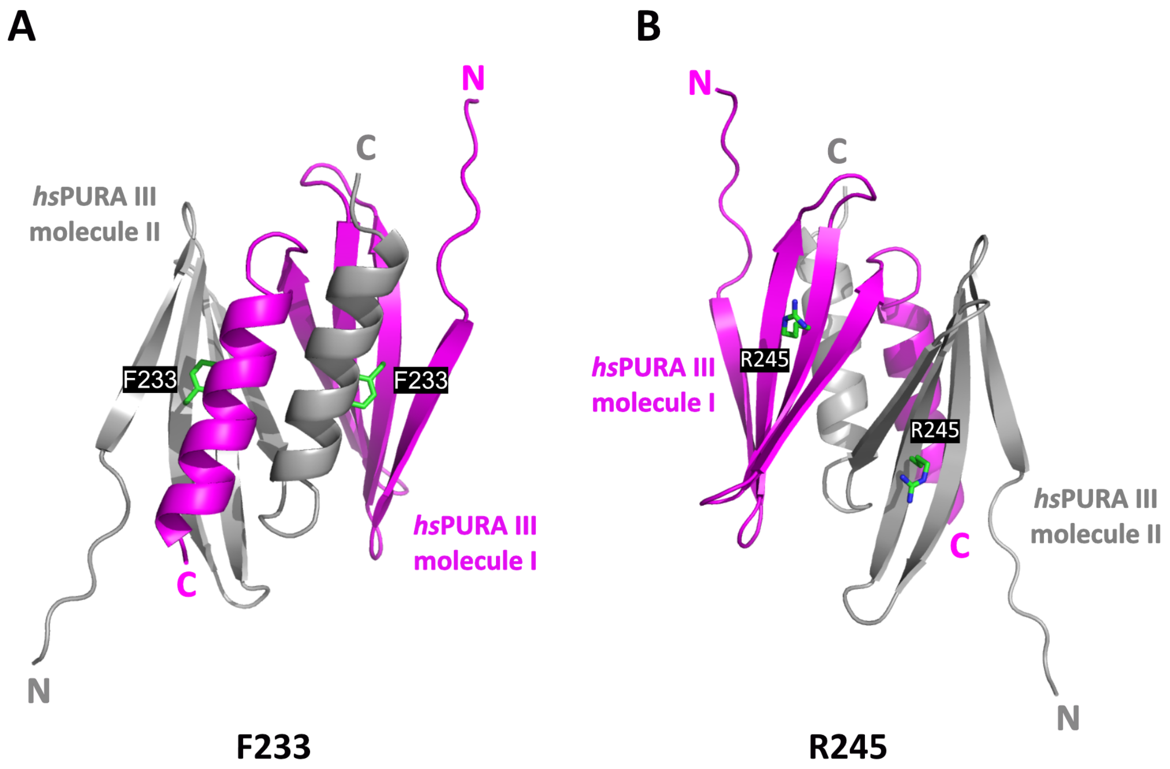


**Appendix Figure S9. The crystal structure of the *hs*PURA III homodimer.**

**A** The crystal structure of the *hs*PURA III homodimer at 1.7 Å resolution (PDB ID: 8CHW) with the residues F233 shown as green sticks and labeled.

**B** The crystal structure of the *hs*PURA III homodimer at 1.7 Å resolution (PDB ID: 8CHW) with the residues R245 shown as green sticks and labeled.

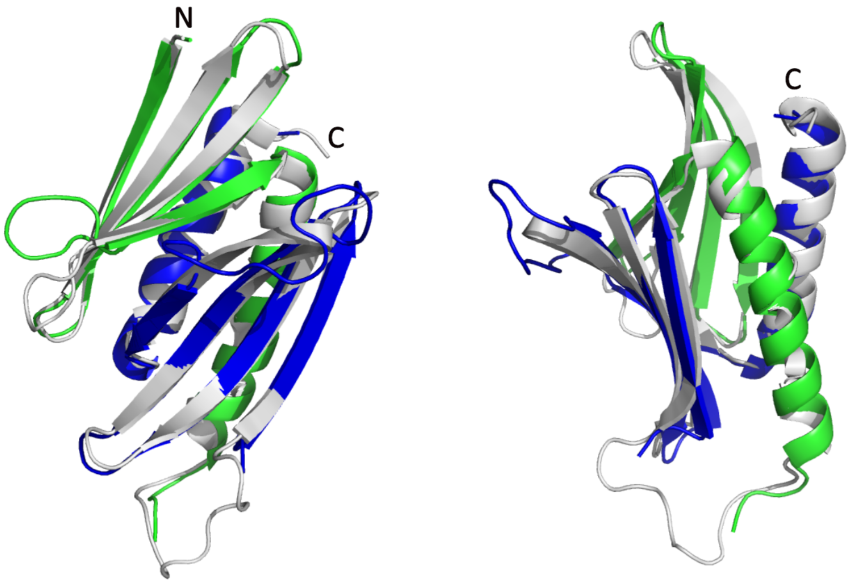


**Appendix Figure S10. Superposition of human and *Drosophila* PURA I-II structure.**

Superposition of *hs*PURA I-II structure (PDB ID: 8CHT; color code as in Figure 4A) with the previously published structure from *Drosophila melanogaster* (*dm*PURA I-II; PDB ID: 3K44; grey ribbon model), with a root mean square deviation (r.m.s.d.) of 0.97 Å for 131 superimposed C$\alpha$ atoms. Superposition is shown in two different orientations.

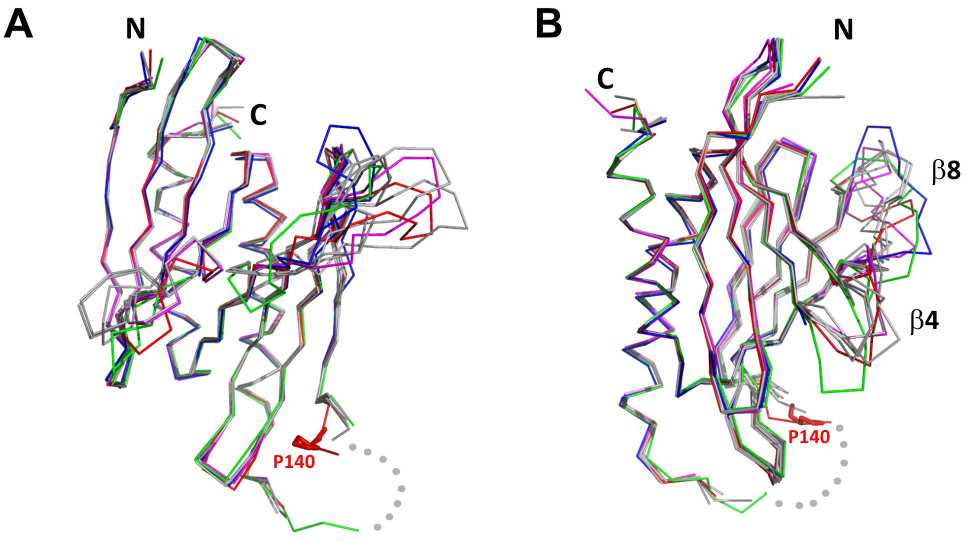


C

| | | *hs*PURA I-II R140P | | | |
|---|---|---|---|---|---|
| | | Chain A | Chain B | Chain C | Chain D |
| *hs*PURA I-II | Chain A | 0.78 / 130 | 0.84 / 135 | 1.00 / 138 | 0.71 / 129 |
| | Chain B | 0.79 / 130 | 0.99 / 136 | 0.92 / 129 | 0.85 / 130 |
| | Chain C | 1.00 / 138 | 1.07 / 134 | 0.82 / 128 | 1.00 / 138 |
| | Chain D | 1.13 / 140 | 0.87 / 132 | 0.80 / 126 | 0.73 / 137 |

**Appendix Figure S11. Structure of *hs*PURA I-II R140P mutant**.

**A-B** An overlay of chains A (red), B (green), C (blue), D (magenta) from the asymmetric unit of the crystals of *hs*PURA I-II R140P, and the wild-type *hs*PURA I-II chains A, B, C, D shown in grey. The structures are shown in two different orientations as ribbons, residue P140 is shown as red sticks. The missing fragment of the loop is shown as grey dots. The most variable part of the secondary structure elements, strands □4 and □8, have been labelled accordingly.

**C** Comparison of all independent chains in the asymmetric unit of *hs*PURA I-II R140P structure (PDB ID: 8CHV) with the wild-type *hs*PURA I-II (PDB ID: 8CHT). R.m.s.d. values are expressed in Å; the number of superimposed C□ atoms is indicated.

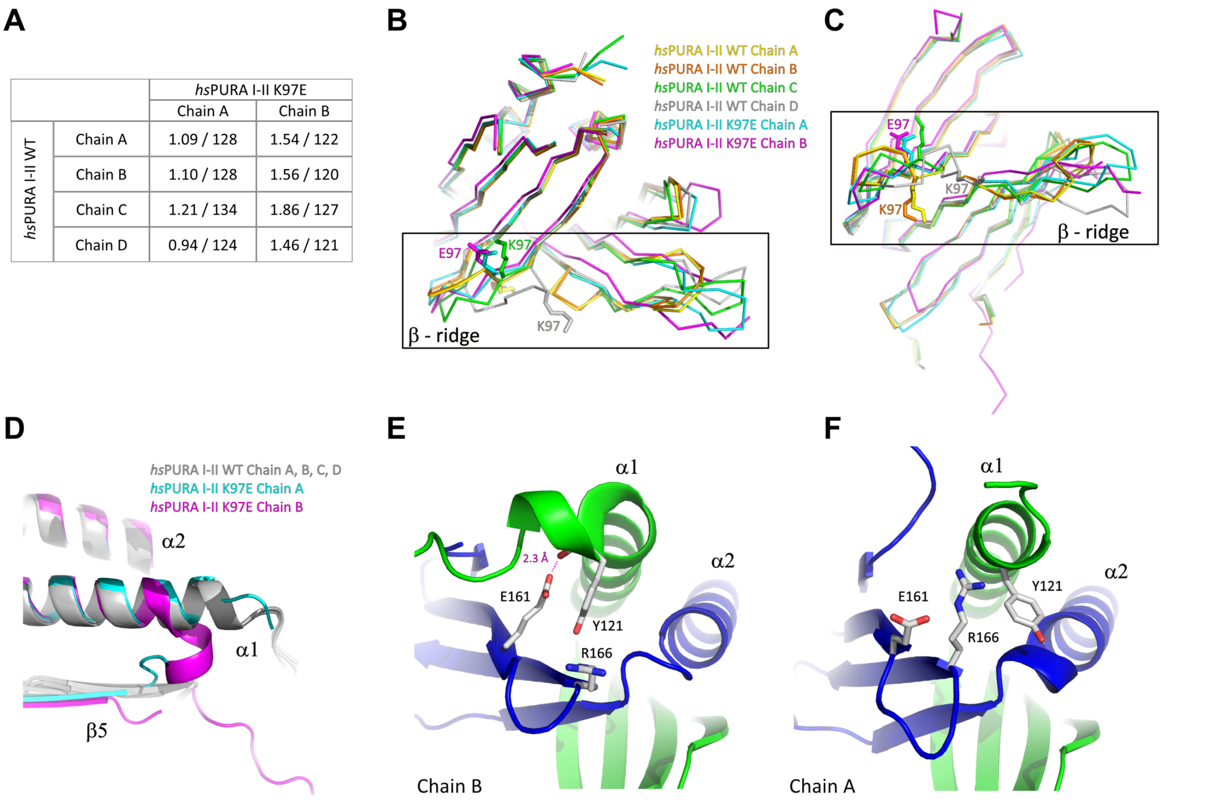


**Appendix Figure S12**. **Comparison of structural features between wild-type and K97E-mutant *hs*PURA protein**.

**A** Comparison of all independent chains in the asymmetric unit of *hs*PURA I-II WT structure (PDB ID: 8CHT) with *hs*PURA I-II K97E (PDB ID: 8CHU). R.m.s.d. values are shown in Å; the number of superimposed Cα atoms has been indicated.

**B-C** The β-ridge in the *hs*PURA I-II crystal structure. The superposition of all four human *hs*PURA I-II chains from the asymmetric unit of the wild-type structure as well as the two chains from the asymmetric unit of the K97E mutant. The structure is shown as ribbons in two different orientations, K97 in the wild-type structure and E97 in the mutated form is shown as sticks.

**D** The overlay of crystal structures of *hs*PURA I-II shown in grey (chains A, B, C, D), *hs*PURA I-II K97E chain A shown in cyan, and *hs*PURA I-II K97E chain B shown in magenta.

**E**-**F** Comparison of the interaction of helix □1 in the chain B (**E**) and the chain A (**F**) in the crystal structure of *hs*PURA I-II K97E. The color code is repeat I: green, repeat II: blue. In chain B the helix □1 bends and forms a new interaction between Y121 and E161 with 2.3 Å distance. This interaction is not present in chain A.

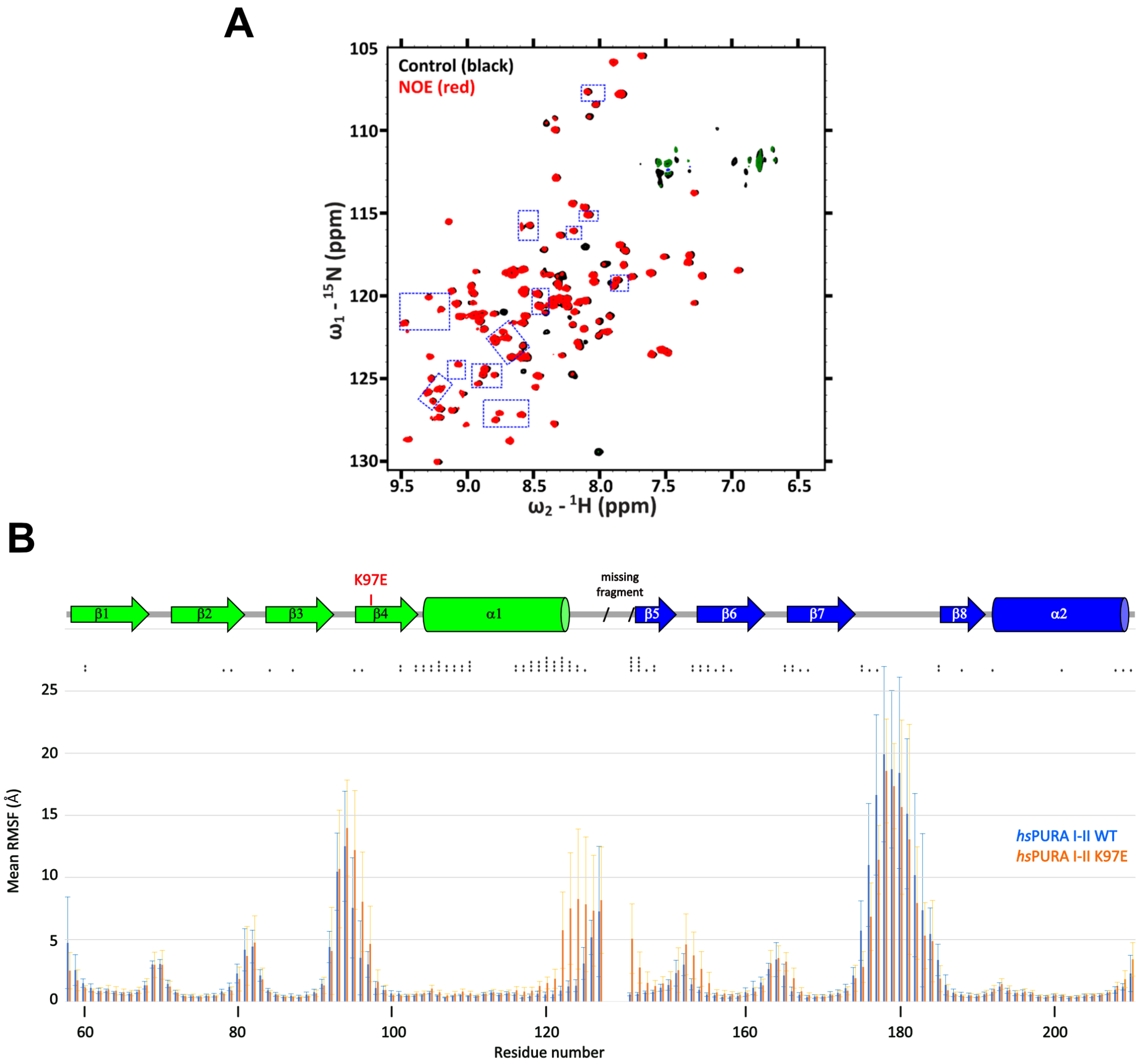


**Appendix Figure S13. Comparison of structural features between wild-type and K97E mutant *hs*PURA protein**.

**A** $^{15}$N-heteronuclear NOE experimental spectra for *hs*PURA I-II K97E, with blue frames indicating changes shown in Fig. 5F. These peaks do not show any shifts compared to the WT control (black), which has no added NOEs, whereas in the experiment with K97E mutant protein (red) additional NOEs are observed.

**B** Summary graph of molecular dynamics simulation for *hs*PURA I-II (blue) and *hs*PURA I-II K97E mutant (orange). The graph shows root mean-square fluctuation in Å (RMSF) for every residue (carbon alpha) of the investigated fragment. RMSF was computed using the average position of the carbon alpha during the simulation as the reference point. Asterisks indicate significance level: * for $p \leq 0.05$, ** for $p \leq 0.01$, *** for $p \leq 0.001$, and **** for $p \leq 0.0001$.

| Protein construct | Purification | Structure | EMSA | dsDNA Strand separation | NanoBRET | Stress Granules | P-bodies |
|---|---|---|---|---|---|---|---|
| PURA I-II WT | ✓ | ✓ | ✓ | ✓ | ✓ | × | × |
| PURA I-II K97E | ✓ | ✓ | ✓ | ✓ | × | × | × |
| PURA I-II R140P | ✓ | ✓ | ✓ | ✓ | × | × | × |
| PURA I-II R199P | × | × | × | × | × | × | × |
| PURA I-II I206F | × | × | × | × | × | × | × |
| PURA I-II m11 | ✓ | × | ✓ | ✓ | × | × | × |
| PURA III WT | ✓ | ✓ | ✓ | ✓ | × | × | × |
| PURA III F233del | × | × | × | × | × | × | × |
| PURA III R245P | × | × | × | × | × | × | × |
| PURA I-III WT | × | × | × | × | ✓ | × | × |
| PURA I-III F233del | × | × | × | × | ✓ | × | × |
| PURA I-III R245P | × | × | × | × | ✓ | × | × |
| PURA FL WT | ✓ | × | ✓ | ✓ | × | ✓ | ✓ |
| PURA FL K97E | × | × | × | × | × | ✓ | ✓ |
| PURA FL I206F | × | × | × | × | × | ✓ | ✓ |
| PURA FL F233del | × | × | × | × | × | ✓ | ✓ |
| PURA FL m11 | × | × | × | × | × | ✓ | ✓ |

**Appendix Table S1.** Summary of the experiments in this study for the selected human PURA variants.

| Data collection | | | | |
|---|---|---|---|---|
| Protein | PURA I-II | PURA I-II K97E | PURA I-II R140P | PURA III |
| PDB ID | 8CHT | 8CHU | 8CHV | 8CHW |
| Beamline | PETRA III DESY P11 | SLS PXIII X06DA | SLS PXIII X06DA | SLS PXIII X06DA |
| Wavelength | 1.033100 | 1.000029 | 1.000040 | 1.000040 |
| Space group | $P2_1$ | $I2_12_12_1$ | $P2_1$ | $P2_12_12$ |
| Cell dimensions<br>*a*, *b*, *c* (Å), □ (°) | 64.92, 58.14, 81.94,<br>□ = 100.65 | 40.02, 77.32, 225.38 | 63.31, 57.58, 84.89, □ = 102.33 | 56.76, 75.64, 31.39 |
| No. of molecules per AU | 4 | 2 | 4 | 2 |
| Resolution (Å) | 50-1.95 (2.00-1.95) | 50-2.45 (2.52-2.45) | 50-2.15 (2.28-2.15) | 50-1.70 (1.74-1.70) |
| $I / \sigma I$ | 12.1 (2.2) | 12.1 (1.28) | 7.6 (0.9) | 19.5 (2.4) |
| CC (1/2) | 99.8 (74.3) | 99.8 (61.9) | 99.7 (58.3) | 100 (88.7) |
| Completeness (%) | 98.0 (96.9) | 99.9 (99.5) | 99.4 (97.7) | 100 (99.9) |
| Redundancy | 6.9 (7.0) | 6.5 (6.6) | 3.5 (3.5) | 6.4 (6.7) |
| **Refinement** | | | | |
| Resolution (Å) | 50-1.95 | 39.43-2.45 | 50-2.15 | 37.85-1.7 |
| No. reflections | 43,138 | 13,362 | 32,410 | 15,494 |
| $R_{work}$ / $R_{free}$ (%) | 17.4/23.1 | 20.2/27.2 | 20.9/28.2 | 17.6/22.4 |
| No. atoms | | | | |
| Protein | 4,659 | 2,287 | 4,597 | 1,108 |
| Water | 314 | 87 | 197 | 132 |
| Other | 16 | 5 | 0 | 34 |
| *B*-factor overall | 26.2 | 61.0 | 52 | 26.4 |
| R.m.s. deviations | | | | |
| Bond lengths (Å) | 0.025 | 0.009 | 0.006 | 0.010 |

| | | | | |
|---|---|---|---|---|
| Bond angles (°) | 2.31 | 1.65 | 1.46 | 1.57 |
| Ramachandran plot | | | | |
| Most favored (%) | 97 | 96 | 95 | 96 |
| Additional allowed (%) | 3 | 4 | 5 | 4 |

**Appendix Table S2.** Data collection and refinement statistics for crystal structures of human PURA variants. Values in parentheses are for the highest-resolution shell.

**Appendix Movie S1.**

Molecular dynamics simulations for the wild-type *hs*PURA I-II (chain B). Red color indicates the residues with significantly higher fluctuation described as p-value computed using the average position of the carbon alpha during the simulation as the reference point. 10 production runs were performed at the temperature of 310K, resulting in 20 simulations of 100 ns each (1000 frames), or a total aggregate time of 1 μs for each protein. The systems were analysed using root means square fluctuations (RMSF). More precisely, the RMSF was computed for each C□ atom in each simulation, leaving 10 RMSF data points per residue.

**Appendix Movie S2.**

Molecular dynamics simulations for the *hs*PURA I-II K97E (chain B). Red color indicates the residues with significantly higher fluctuation described as p-value computed using the average position of the carbon alpha during the simulation as the reference point. 10 production runs were performed at the temperature of 310K, resulting in 20 simulations of 100 ns each (1000 frames), or a total aggregate time of 1 μs for each protein. The systems were analysed using root means square fluctuations (RMSF). More precisely, the RMSF was computed for each C□ atom in each simulation, leaving 10 RMSF data points per residue.

### MATERIALS AND METHODS

#### Establishment of stable cell lines

HeLa Kyoto cells were grown in DMEM (4.5 g/l glucose, L-glutamine, phenol red; Invitrogen), supplemented with 10% FBS, and 100 U/ml Penicillin/Streptomycin. Stable cell lines were established using the PiggyBac transposon system and transfected with Lipofectamine 3000. After three days of incubation, the medium was exchanged and supplemented with 700 µg/ml Hygromycin B (Invitrogen) for selection over three weeks. Stable cell lines were sorted for GFP signal by flow cytometry for similar expression levels at the core facility Flow Cytometry of the Biomedical Centre Munich (Munich University). Expression levels were confirmed by Western blotting.

#### Stress treatment

For each cell line, protein expression in 1 x $10^5$ cells was induced by doxycycline at a final concentration of 1 µg/ml the day before stress treatment. Cells were treated for one hour with 500 µM sodium arsenite to induce stress-granule formation. As a reference, untreated cells received only media. After stress induction, stress media was removed, and cells were analyzed by immunofluorescence (IF) staining.

#### Immunofluorescence staining

Cells grown on NO 1.5 coverslips were washed once with PBS, fixed with 3.7% formaldehyde (diluted in PBS) for 10 min, and washed twice with PBS. Then, cells were incubated for 10 min with blocking buffer (1% donkey serum diluted in PBST), before adding the primary antibody (diluted in blocking buffer) for one hour at RT. Subsequently, cells were washed three times for 5 min with PBST and the secondary antibody (diluted in blocking buffer) was added for 1 h at RT in the dark. Afterward, cells were washed three times with PBST before DAPI (0.5 µg/ml) staining and mounting of the coverslips in Prolong Diamond Antifade Mountant onto the microscope slide.

**Microscopy**

Confocal microscopy was performed at the Biomedical Center (Core facility Bioimaging; Munich University) with an inverted Leica SP8 microscope, equipped with lasers for 405, 488, 552, and 638 nm excitation. Either a 100 x 1.4 or a 63 x 1.4 oil objective was used to acquire the images (pixel size: 80 nm) using the fluorescence settings DAPI: 415-470 nm, GFP: 498-535, Cy3: 562-620, Cy5:648-710. GFP, Cy3, and Cy5 were recorded with hybrid photodetectors (HyDs), DAPI with a conventional photomultiplier tube.

**Quantitative analysis of microscopic images with ImageJ**

Images were processed using the Fiji:ImageJ software, applying only linear enhancements for brightness and contrast. Within one experiment, equal exposure times and processing conditions were used for all samples in respective channels. Co-localization of *hs*PURA with DCP1A and G3BP1 was quantified using the magic wand tool of Fiji:ImageJ software to determine the intensities ($I_{granule}$) of the processing bodies (PBs) or stress granules. Then, a margin of 0.5 µm for PBs and 0.3 µm for stress granules was drawn around the granule structure and intensities measured ($I_{band}$). Background intensity ($I_{background}$) was determined at a spot outside of cells and finally, the ratio of $I = \frac{I_{granule} - I_{background}}{I_{band} - I_{background}}$ was calculated.

**siRNA-mediated *PURA* knock-down in HeLa cells**

The siRNA-mediated PURA knock-down was performed as described previously (Molitor *et al*, 2023). Briefly, approximately 1 x $10^5$ HeLa cells were plated in 24-well plates- The next day, when the cells reached around 30% confluency, lipofection was performed using RNAiMAX (ThermoFisher Scientific) according to the manufacturer´s instruction. For that, a pre-designed PURA siRNA pool was used (Dharmacon, M-012136-01-000) and as a control siGENOME Non-Targeting Pool #1 (Dharmacon, D-001206-13-05). 20 µm of siRNA, either PURA siRNA or Control siRNA, was mixed with OptiMEM (ThermoFisher Scientific) in one reaction tube, and lipofectamine RNAiMAX was mixed with OptiMEM in another. After an incubation time of 5-10 min at room temperature, both reactions were mixed and further incubated for 20 min at room temperature. Subsequently, the culture media of HeLa cells was changed to 250 µL of FMEM +10% FBS + 1% Penicillin/Strepatvidin. Finally, 100 µL of the lipofection mixture was added drop-wise on each 24-well of HeLa cells. The transfected cells were incubated for 72 hours at 37°C and 5% $CO_2$ before being further used for Immunofluorescence analysis.

#### Quantitative analysis of immunofluorescence staining in *PURA*-knockdown experiments

Quantification of stress granules in PURA wild-type versus PURA knockdown HeLa cells was done using Fiji software (version 2.3.0/1.53q) (Molitor *et al.*, 2023). Zeiss files (czi) were opened in Fiji and the Channels were split in DAPI and G3BP1 channels. Then, the cell number was determined by using the auto threshold function on the DAPI channel and analyzing particles (Analyze particle function) with a particle size of 1000-infinity pixel units, which should cover all DAPI signals. G3BP1-granules were quantified using the same functions but selecting a particle size of ≥ 0.7 microns$^2$ and a circularity of 0.1-1.00. Size comparison was done by comparing the average size per granule within the particle size analysis function. Quantification of stress granules was done in biological triplicates and technical triplicates with a magnification of 20x.

#### Purification of total HeLa RNA

RNA from 1x$10^7$ HeLa cells was extracted using the TRIzol Plus RNA Purification Kit (Thermo Fisher Scientific) according to the manufacturer's instructions and following the general precautions required for RNA work. RNA was eluted in 100 □l DEPC water. Integrity, purity, and amount of total HeLa RNA were determined using Agilent Tape station measurements according to the manufacturer's instructions.

#### *In vitro* phase separation assays

Full-length wild-type *hs*PURA (aa 1-322) and FUS RGG3-PY (aa 454-526) were thawed and FUS RGG3-PY was incubated at 95°C for 5 min and subsequently kept at room temperature. Proteins were diluted in condensate buffer (20 mM $Na_2HPO_4$/$NaH_2PO_4$, pH 7.5, 150 mM NaCl, 2.5% glycerol, 1 mM DTT), and total HeLa cell RNA was added at an RNA-to-protein mass ratio of 0.5. For visualization of condensates, samples were immediately transferred to self-assembled sample chambers formed by double-sided sticky tape, taped onto a glass slide, and sealed with a coverslip. Condensates were imaged after 15 min using phase contrast microscopy and a HC PL Fluotar L 40x/0.6 PH2 objective on a Leica DMi8 microscope (Leica, Germany).

### NanoBRET

HEK293 cells were seeded in 6-well plates at 800,000 cells per well with culture media (89% DMEM (Gibco); 10% Fetal Bovine Serum (FBS); 1% Anti-Anti (Gibco)) and incubated for 6 h at 37°C / 5% $CO_2$. Cells were transfected using Lipofectamine 3000 (Thermo Fisher Scientific) and a ratio of 1:100 for HaloPURAI-III and NLucPURAI-III mutants. Cells were incubated for 24 h at 37°C / 5% $CO_2$, trypsinized, adjusted to a final density of 200,000 cells/ml with assay medium (95% Opti-MEM® I Reduced Serum Medium; 4% FBS; 1% Anti-Anti), and divided into two pools. Next, either HaloTag®NanoBRET 618 Ligand or dimethyl sulfoxide (DMSO) as no-ligand control was added to each pool at concentrations of 0.1 M and 0.1%, respectively. Three technical replicates of each cell suspension (20,000 cells) were transferred in a white Lumitrac 96-Well plate (F-Bottom) and incubated for 18 h at 37°C/5% $CO_2$. 25 µl diluted NanoBRET™ Nano-Glo Substrate (1:100 in Opti-MEM® I Reduced Serum Medium, no phenol red) was added to all wells and mixed for 30 seconds. Plates were measured using a Tecan Spark plate reader with the following settings: HaloTag®NanoBRET 618 Ligand: 595 nm – 650 nm (Bandwidth 27.5 nm); NanoLuciferase®: 430 nm- 455 nm (Bandwidth 12.5 nm); Integration time: 1,000 ms. Samples were provided as biological triplicates, of which each was measured as technical duplicates. As positive control the homotypic dimerization of wild-type *hs*PURA I-III was applied as a single sample on each 96-well plates. The experimental mBUs (milliBRET units) were calculated by (618 EM / 460 EM) x 1,000 x BU = mBU. From these mBU the no-ligand control was subtracted.

### Cloning, expression, and purification of *hs*PURA repeat I-II

*hs*PURA protein fragment E57-E212 (PUR repeat I and II) was amplified by PCR and cloned using in-fusion method (Berrow *et al*, 2007) into pOPINS3C vector and expressed in *E. coli* Rosetta cells using auto-induction media (Studier, 2005). The cell pellet was lysed in a buffer composed of 50 mM HEPES pH 7.5, 500 mM NaCl, 20 mM imidazole, supplemented with protease inhibitors (Roche) and DNAse, and centrifuged. The supernatant was applied on a 5 mL HisTrap column (GE Healthcare), washed with high salt buffer (50 mM HEPES pH 7.5, 1 M NaCl,

20 mM imidazole), and eluted with a linear gradient from 20 to 400 mM imidazole. Protein was dialyzed against 50 mM HEPES 7.5, 200 mM NaCl, 1 mM DTT overnight at 4°C with PreScission protease added. After a subtractive HisTrap column, the flow-through was diluted in 50 mM HEPES pH 7.5, 70 mM NaCl and loaded onto a 1 mL HiTrap Heparin HP (GE Healthcare) column and eluted using a linear NaCl gradient (0.1-2 M) and subsequently applied onto a Superdex75 16/60 column (GE Healthcare) equilibrated with 20 mM HEPES pH 7.5 and 200 mM NaCl. The selected fractions were validated via SDS PAGE, pooled, and concentrated to 8 mg/ml.

**Cloning, expression, and purification of *hs*PURA K97E repeat I-II**

*hs*PURA protein fragment E57-E212 with K97E mutation was cloned into pOPINJ expression vector as fusion protein with N-terminal His$_6$-GST-tag. After expression in *E. coli* Rosetta cells with LB media at 18°C and induction with IPTG (0.25 mM) at OD 0.6, cells were lysed by sonication in resuspension buffer (1 M NaCl, 50 mM HEPES, 2 mM DTT, pH 7.5, DNAse I, EDTA free protease inhibitor (Roche)) and centrifuged for 30 min at 20,000 x g at 4°C. The soluble protein fraction was bound to a GSTrap FF column (GE Healthcare) and washed with lysis buffer, high salt buffer (2 M NaCl, 20 mM HEPES, 2 mM DTT, pH 7.5), and dialysis buffer (300 mM NaCl, 20 mM HEPES, 2 mM DTT, pH 7.5 at 4°C). Bound protein was eluted with buffer containing 500 mM NaCl, 20 mM HEPES, 25 mM Glutathione, pH 7.5. Protein was dialyzed overnight in dialysis buffer together with PreScission protease (100 □g per 5 ml). After a subtractive GSTrap column, the flow-through was diluted in 50 mM HEPES pH 7.5, 70 mM NaCl and loaded onto a 5 mL HiTrap Q FF (GE Healthcare) column and eluted using a linear NaCl gradient (0.1-2 M). The same procedure was repeated using HiTrap Heparin HP (GE Healthcare) column. As the last step, size exclusion chromatography has been performed in buffer containing 300 mM NaCl, 20 mM HEPES, 2 mM DTT, pH 7.5 at 4°C on Superdex75 16/60 column (GE Healthcare). The selected fractions were validated via SDS PAGE, pooled, and concentrated to 8.2 mg/ml.

**Cloning, expression, and purification of *hs*PURA R140P repeat I-II**

*hs*PURA protein fragment E57-E212 with R140P mutation was cloned into pOPINJ expression vector as fusion protein with N-terminal His$_6$-GST-tag and expressed in *E. coli* Rosetta cells using auto-induction media at 22°C. The purification has been performed as described for *hs*PURA repeat I-II K97E mutant. The selected fractions were validated via SDS PAGE, pooled, and concentrated to 5.6 mg/ml.

#### Cloning, expression, and purification of *hs*PURA repeat I-II m11 mutant

*hs*PURA protein fragment E57-E212 with 11 mutations abolishing nucleic acid binding (K71A, N80A, K82G, F85A, K87A, K97A, R153A, N162A, R164G, F167A, and R169A) was cloned into pOPINJ expression vector. The purification was performed using the same protocol as for *hs*PURA repeat I-II K97E mutant. The selected fractions were validated via SDS PAGE, pooled, and concentrated to 5.2 mg/ml.

#### Cloning, expression, and purification of full-length WT *hs*PURA and full-length *hs*PURA m11 mutant

The full-length wild-type *hs*PURA as well as its m11-mutant variant were cloned into pOPINJ vector. Expression and purification of both proteins were performed as described above for K97E mutant. The selected fractions were validated via SDS PAGE, pooled, and concentrated to 3.4 mg/ml (wt) and 1.6 mg/ml (m11), respectively.

#### Cloning, expression, and purification of *hs*PURA repeat III

Fragment P215-K280 of *hs*PURA (PUR repeat III) was amplified by PCR and cloned into pOPINS3C vector. Expression and purification were performed as described above for *hs*PURA repeat I-II except for the exclusion of the Heparin column step. The selected fractions were validated via SDS PAGE, pooled, and concentrated to 7.5 mg/ml.

#### Electrophoretic mobility shift assay (EMSA)

Electrophoretic mobility shift assays were performed in a final reaction volume of 20 □L in EMSA buffer composed of 300 mM NaCl, 20 mM HEPES pH 8.0, 2 mM DTT, 4% (v/v) glycerol, and 3 mM $MgCl_2$. The final protein concentration ranged from 0 to 16 □M, with a constant final concentration of fluorescence-labeled RNA 8 nM and 10 □g/mL of competitor yeast tRNA. The

24-mer RNA fragment $(CGG)_8$ was labeled on its 5’ end with Cy5 fluorophore (Eurofins Genomics). After incubation of the reaction for 30 min at RT, 5 µL of the samples were loaded onto the 6% TBE polyacrylamide gel and run for 45 min at 100 V. The fluorescence measurements were performed using an Amersham Typhoon scanner and software v 2.0.0.6. The intensities of the bands observed on the scanned EMSAs gels were quantified with FUJI ImageJ software (v 1.53C).

**Unwinding assay**

For the fluorescence-based unwinding experiment the double-labeled DNA fragment, FAM-**CCAGG**GCACTTAAAAAAATTCG**CCTGG**-Dabcyl, was used (double-stranded part of the DNA is shown as bold). The assay was performed in 50 µL reaction volume in the unwinding buffer composed of 300 mM NaCl, 20 mM HEPES pH 8.0, 2 mM DTT, 4% (v/v) glycerol, and 3 mM $MgCl_2$. The protein concentration ranged from 0 to 16 µM, with a constant DNA concentration of 50 nM. The fluorescence measurements were performed using a Perkin Elmer EnVision 2104 Multilable Reader and Wallac EnVision Manager software v 1.12. FAM fluorescence was excited at 495 nm with a slit of 2 nm. Emission was recorded at 517 nm with a slit of 3 nm for 0.5 s (integration time). Experiments were performed at least three times.

**Circular dichroism**

For secondary structure analysis of *hs*PURA I-II wt, K97E, and R140P, CD spectra were recorded using a Jasco J-715 spectropolarimeter (JASCO) and the range of 190 nm to 260 nm at 20°C. The measurements were performed with a 1 mm path length high precision quartz cuvette (Hellma Analytics). The concentration of all tested *hs*PURA variants was 20 µM in a buffer containing 60 mM NaCl, 20 mM HEPES, and 2 mM DTT at pH 7.5. Further measurement parameters included a scanning speed of 50 nm/min, 3 scans, and a response time of 8 s. Sensitivity was set to standard (100 mdeg).

### Crystallization

The crystallization experiments for *hs*PURA I-II, *hs*PURA I-II K97E, *hs*PURA I-II R140P and *hs*PURA III were performed at the X-ray Crystallography Platform at Helmholtz Munich. For *hs*PURA I-II, the best diffracting crystals were obtained from 0.1 M Tris buffer pH 8.2, 0.2 M sodium acetate, and 30% (w/v) PEG 4,000. The best crystals for *hs*PURA I-II K97E grew in 0.1 M citric acid pH 4.0, 1 M LiCl, 9% (w/v) PEG 6,000. *hs*PURA I-II R140P crystallized directly during protein concentration in the size exclusion buffer. For *hs*PURA III crystals were grown in 0.1 M Tris pH 8.5, 12% (v/v) glycerol, 1.5 M $(NH_4)_2SO_4$. For the X-ray diffraction experiments, crystals were mounted in the nylon fiber loops and flash-cooled to 100 K in liquid nitrogen. The cryoprotection was performed for a few seconds in reservoir solution complemented with 25% (v/v) ethylene glycol in all cases. Diffraction data for *hs*PURA I-II were collected on the PETRAIII P11 beamline, for *hs*PURA I-II K97E, *hs*PURA I-II R140P and *hs*PURA III the measurements were performed at Swiss Light Source, Paul Scherrer Institute, beamline X06DA. The best data set for each protein variant was indexed and integrated using *XDS* (Kabsch, 2010) and scaled using *SCALA* (Evans, 2006; Winn *et al*, 2011). Intensities were converted to structure-factor amplitudes using the program *TRUNCATE* (French & Wilson, 1978). Supplementary Table S1 summarizes data collection, processing, and refinement statistics.

### Structure determination and refinement

The structures of *hs*PURA I-II and *hs*PURA III were solved by molecular replacement using the crystal structures of *Drosophila* PURA repeat I-II (PDB ID: 3k44) (Graebsch *et al*, 2009) and repeat III (PDB-ID: 5FGO) (Weber *et al*, 2016) as search model, respectively. The structures of *hs*PURA I-II K97E and *hs*PURA I-II R140P were solved by molecular replacement using the structure of *hs*PURA I-II from this study as search model. In all cases, for the molecular replacement calculations the program *MolRep* (*CCP4*) was used. Model rebuilding in all cases was performed with *COOT* (Emsley & Cowtan, 2004). The refinement was done with *REFMAC5* (Murshudov *et al*, 1997) using the maximum-likelihood target function. The final model is characterized by R and $R_{free}$ factors of 17.4/23.1%, 20.2/27.2%, 20.9/28.2% and 17.6/22.4% for *hs*PURA I-II, *hs*PURA I-II K97E, *hs*PURA I-II R140P and *hs*PURA III, respectively. The stereochemical analysis of the final model was done with *PROCHECK* (Laskowski *et al*, 1993) and *MolProbity* (Chen *et al*, 2010). For all the crystallographic calculations, the *SBGrid* software bundle was used (Morin *et al*, 2013). Atomic coordinates and structure factors have been deposited in the Protein Data Bank under the accession codes 8CHT (*hs*PURA I-II), 8CHU (*hs*PURA I-II K97E), 8CHV (*hs*PURA I-II R140P) and 8CHW (*hs*PURA III).

**Expression of isotope-labeled proteins for NMR**

$^{15}$N labelled proteins (*hs*PURA I-II wt and K97E mutant) were expressed in $^{15}$N-M9 minimal medium (1□□□$^{15}$N-labeled M9 salt solution, 0.4% glucose, 1 mM $MgSO_4$, 0.3 mM $CaCl_2$, 1 µg /L biotin, 1 µg/L thiamine, 1□ trace metals) with respective antibiotics. 150 ml of pre-culture was grown overnight at 37°C and used for inoculation of 3L pre-warmed M9 minimal medium the next day. After growing to $OD_{600\ nm}$ = 0.6 at 37°C, cells were induced with 0.25 mM IPTG, and proteins expressed at 18°C overnight (O/N) (approx. 18 h).

**NMR**

NMR measurements for HSQC comparison were performed with the $^{15}$N labeled samples in buffer containing 300 mM NaCl, 20 mM HEPES, 0.2 mM TCEP, and pH 7.5. Uniformly $^{15}$N-labeled NMR samples were prepared at protein concentrations of 25 µM (WT and K97E mutant) and 454 µM (K97E) in a buffer containing 300 mM NaCl, 20 mM HEPES, 0.2 mM TCEP, and pH 7.5 with 10% $D_2O$ for lock signal. NMR experiments were recorded at 298 K on 800- and 600-MHz Bruker Avance NMR spectrometers, equipped with cryogenic or room-temperature triple resonance gradient probes. NMR spectra were processed by *TOPSPIN3.5* (Bruker), then analyzed using *NMRFAM-SPARKY* (Lee *et al*, 2015). 1H-$^{15}$N Heteronuclear NOE experiments were recorded on a 600-MHz spectrometer at 298 K with an interleaved manner with and without proton saturation.

**Molecular dynamics (MD) simulations**

Molecular dynamics simulations were performed using the *ACEMD* engine (Harvey *et al*, 2009). The system was prepared for simulation using the HTMD library (Doerr & De Fabritiis, 2014). Both, the *hs*PURA I-II WT and the K97E mutant, were prepared starting from the available structures (PDB ID: 7PO8 - WT, chain B; and PDB ID: 7PRF - K97E mutant, chain B). Due to lack of resolved structure of the loop between □1 and □5 (the electron density map was not visible) for the MD calculations the missing parts were manually adjusted and linked. This part of the structure was not taken for the final analysis. The proteins were titrated using HTMD Protein Prepare, adding hydrogens and generating ionization states for the side chains using PROPKA at pH 7.0, and end-capping of N- and C-termini, using ACE and NME capping. The prepared proteins were parameterized using the Amber 14SB force field (Wang *et al*, 2004). The systems were solvated in a box of dimensions 70.6, 70.6, 70.6 Å for the K97E and 67.1, 67.1, 67.1 for WT; the chosen solvent model was TIP3P. The solvated systems were equilibrated for 3 ns using HTMD equilibration protocols applying restraints on protein backbones for minimization. Equilibration was run at 310K with NPT conditions setting the cut-off for long-range PME forces to 9 Å; a time step of 4 fs was used for the integration and energy calculation. 10 production runs were performed on the resulting equilibrated system at the temperature of 310K for WT and K97E, resulting in 20 simulations of 100 ns each (1000 frames), or a total aggregate time of 1 μs for each protein. The systems were analysed using root means square fluctuations (RMSF). More precisely, the RMSF was computed for each C□ atom in each simulation, leaving 10 RMSF data points per residue. The mean and standard deviation of these data points were calculated, and a two-sided t-student test was performed on each pair of comparable residues, that is, residues that are paired in the sequence alignment of the two proteins.
